# Short-term changes in polysaccharide utilization mechanisms of marine bacterioplankton during a spring phytoplankton bloom

**DOI:** 10.1101/2020.01.08.899153

**Authors:** Greta Reintjes, Bernhard M. Fuchs, Mirco Scharfe, Karen H. Wiltshire, Rudolf Amann, Carol Arnosti

## Abstract

Spring phytoplankton blooms in temperate environments contribute disproportionately to global marine productivity. Bloom-derived organic matter, much of it occurring as polysaccharides, fuels biogeochemical cycles driven by interacting autotrophic and heterotrophic communities. We tracked changes in the mode of polysaccharide utilization by heterotrophic bacteria during the course of a diatom-dominated bloom in the German Bight, North Sea. Polysaccharides can be taken up in a ‘selfish’ mode, where initial hydrolysis is coupled to transport into the periplasm, such that little to no low molecular weight (LMW) products are externally released to the environment. Alternatively, polysaccharides hydrolyzed by cell-surface attached or free extracellular enzymes (external hydrolysis) yield LMW products available to the wider bacterioplankton community. In the early bloom phase, selfish activity was accompanied by low extracellular hydrolysis rates of a few polysaccharides. As the bloom progressed, selfish uptake increased markedly, and external hydrolysis rates increased, but only for a limited range of substrates. The late bloom phase was characterized by high external hydrolysis rates of a broad range of polysaccharides, and reduced selfish uptake of polysaccharides, except for laminarin. Substrate utilization mode is related both to substrate structural complexity and to the bloom-stage dependent composition of the heterotrophic bacterial community.

**Originality statement:** The means by which heterotrophic bacteria cooperate and compete to obtain substrates is a key factor determining the rate and location at which organic matter is cycled in the ocean. Much of this organic matter is high molecular weight (HMW), and must be enzymatically hydrolyzed to smaller pieces to be processed by bacterial communities. Some of these enzyme-producing bacteria are ‘selfish’, processing HMW organic matter without releasing low molecular weight (LMW) products to the environment. Other bacteria hydrolyze HMW substrates in a manner that releases LMW products to the wider bacterial community. How these mechanisms of substrate hydrolysis work against a changing background of organic matter supply is unclear. Here, we measured changing rates and mechanisms of substrate processing during the course of a natural phytoplankton bloom in the North Sea. Selfish bacteria generally dominate in the initial bloom stages, but a greater supply of increasingly complex substrates in later bloom stages leads to external hydrolysis of a wider range of substrates, increasing the supply of LMW hydrolysis products to the wider bacterial community.

## Introduction

Spring phytoplankton blooms in temperate environments are responsible for a considerable portion of annual primary productivity, and via their production and demise, are coupled closely to the transformation and remineralization of organic matter throughout the oceanic food chain. Although the specific interactions among phytoplankton, nutrients, water column mixing, and grazers mean that spatial extents and temporal courses of spring blooms are highly variable (Martinez *et al*., 2011; Daniels *et al*., 2015), dynamic changes in primary producer communities during spring blooms can be closely tracked (Wiltshire *et al*., 2015; Sarker *et al*. 2018; Scharfe and Wiltshire, 2019). Less is known, however, about the composition and structure of complex organic matter that they produce in response to nutrient and light availability, and as they are affected by viruses and grazing (e.g. Daniels *et al*., 2015).

This complex organic matter is a direct link coupling phytoplankton and the heterotrophic bacterial communities that are responsible for recycling a substantial fraction of primary production (Teeling *et al*., 2012; Buchan *et al*., 2014). The bacterial response to rapidly increasing concentrations of phytoplankton-derived organic matter during blooms is typically dominated by ‘boom-and-bust’ specialists who opportunistically react to rapidly changing conditions, including members of the *Bacteroidetes*, *Gammaproteobacteria*, and *Roseobacter* (*Alphaproteobacteria*) (Buchan *et al*., 2014). Within these taxa, successive bacterial groups grow and replace one another (Teeling *et al*., 2012), with a marked temporal succession in key genes and proteins produced by individual bacterial groups that are linked with the processing of complex organic matter (Teeling *et al*., 2012; 2016). The nature and structures of the phytoplankton-derived substrates targeted by specific bacterial groups is only beginning to be revealed through focused genetic and (meta-)genomic analyses that indicate the substrate specialization of specific organisms (Teeling *et al*., 2012; 2016; Kappelmann *et al*., 2018; Krüger *et al*., 2019).

Measurements of the rates at which these bacteria degrade their target substrates, and the turnover rates of complex organic matter pools during a phytoplankton bloom are still lacking, however. Here we seek to fill this gap by measuring hydrolysis rates of polysaccharides, an abundant class of high molecular weight organic matter in algae (Painter, 1983), and simultaneously tracking changes in the bacterial community during the course of a phytoplankton bloom at Helgoland (North Sea). Polysaccharides must be enzymatically hydrolyzed to smaller sizes in order to be used as substrates by bacterial communities. The structural specificities of the extracellular enzymes produced by bacteria thus help determine which polysaccharides are available for metabolism by the wider microbial community (Arnosti, 2011). Over a six-week period during the 2016 bloom, we followed basic biological (bacterial cell counts, phytoplankton abundance) and physical/chemical parameters (N, P, Si, Temp). At four distinct timepoints, we measured the rates and mechanisms by which bloom-associated bacterial communities processed complex polysaccharides.

Enzymatic processing of polysaccharides by bacteria in the ocean has recently been discovered to be carried out by two distinctly different mechanisms: the first mechanism, long known in microbiology, is referred to here as “external hydrolysis”: polysaccharides are hydrolyzed to low molecular weights outside the cell by cell-surface-associated or free extracellular enzymes. The low molecular weight products of external hydrolysis can be taken up by a range of bacteria, including those that did not produce the extracellular enzymes. A second mechanism, “selfish” substrate processing (Cuskin *et al.,* 2015), builds on a polysaccharide degradation mechanism known for more than a decade from gut microbiology (Cho and Salyers, 2001; Martens *et al.,* 2009). With ‘selfish’ uptake, initial hydrolysis of a polysaccharide is coupled directly to transport of the resulting large oligosaccharides into the periplasmic space, producing little or no low molecular weight hydrolysis products in the external environment (Cuskin *et al*. 2015; Rakoff-Nahoum *et al*. 2016). Only recently has the existence and prevalence of ‘selfish’ uptake been recognized in surface waters of the Atlantic Ocean (Reintjes et al., 2017; 2019). Across a transect of the North and South Atlantic, the relative contributions of selfish uptake and external hydrolysis to polysaccharide degradation varied by substrate, as well as by location (Reintjes *et al*., 2019), with selfish uptake reaching up to 25% of DAPI-stainable cells (Reintjes *et al*., 2017).

We hypothesize that a range of factors, including substrate availability and complexity, controls the extent of selfish and external hydrolysis at a given location (Arnosti *et al*., 2018). Our previous investigations (Reintjes et al., 2017; 2019), however, were made at single timepoints across a broad transect; in effect, we had point measurements of bacterial communities whose past histories we could only infer. The current investigation provided the opportunity to track both mechanisms of polysaccharide degradation against a clearly-defined background of changing phytoplankton communities, which constitute the major source of substrates for heterotrophic bacteria.

Here, we tracked dynamics in phytoplankton and bacterial communities as they responded to changing conditions over the course of a spring bloom. Our focus was particularly on the prevalence and presence of selfish uptake and external hydrolysis over the course of the bloom. We measured both processes by incubating fluorescently-labeled polysaccharides (FLA-PS; Arnosti, 2003) in water samples collected at four timepoints during different bloom phases. Time-course subsamples were collected from each incubation to monitor the extent of external hydrolysis, as well as selfish uptake of large fragments of FLA-PS. Samples were also collected for next generation sequencing (NGS) to monitor changes in community composition. External hydrolysis of the FLA-PS was measured as the systematic decreases with time of the molecular weight of the added polysaccharide in the incubation water, a process that can be tracked analytically via gel permeation chromatography and fluorescence detection (Arnosti, 2003). Selfish uptake of the polysaccharides was visualized microscopically in subsamples of water as uptake of the glycan into the periplasmic space of bacteria (Reintjes *et al.,* 2017). Fluorescent staining in the periplasmic space, particularly at time points in an incubation at which no low molecular weight hydrolysis products are detectable in the external environment, is a clear indication of selfish uptake (Reintjes *et al.,* 2017). Monitoring the size distribution of FLA-PS in the external environment, as well as the presence of fluorescence in the periplasmic space of bacteria, thus enables us to track in the same incubations mechanisms of polysaccharide processing, as well as rates of enzymatic hydrolysis.

Understanding the controls on external hydrolysis and selfish uptake can provide new insight into factors affecting microbially-driven carbon cycling and the composition of heterotrophic microbial communities in surface ocean waters. The mechanism by which high molecular weight polysaccharides are processed in the environment – external hydrolysis, or selfish uptake – can affect the size distribution of bloom-derived organic matter in the water column and thus the extent to which low molecular weight (LMW) hydrolysis products are made available to the rest of the microbial community (Arnosti *et al.,* 2018). The pool size of LMW hydrolysis products available can in turn affect population dynamics of “scavenging” bacteria that take up LMW hydrolysis products, although they do not produce extracellular enzymes (Arnosti *et al.,* 2018; Reintjes *et al.,* 2019). The relative balance of selfish uptake and external hydrolysis is thus relevant to the workings of the marine carbon cycle and dynamics of heterotrophic microbial communities.

## Results

### Bloom development: nutrients, diatoms, and bacterial counts and composition

A spring bloom developed in 2016 consistent with previous observations carried out using continuous measurements at Helgoland Roads (https://doi.pangaea.de/10.1594/PANGAEA.864676; Fig. S1). Temperature increased steadily from 6.2 °C to 8.8 °C between our sampling timepoints on March 22^nd^, April 5^th^, April 19^th^, and May 3^rd^, 2016. These sampling dates are referred to as Hel_1 to Hel_4 in the following text. Nutrient concentrations indicated the development of the bloom, with silicate decreasing from 8.47 μM for Hel_1 to concentrations close to 0.5 μM for Hel_2 to Hel_3, and 0.13 μM for Hel_4. Nitrate concentrations over the same time points also decreased sharply, with concentrations of 21.4 μM on Hel_1 to 9.4, 7.2 and 7.7 μM for Hel_2, Hel_3, and Hel_4. Phosphate concentrations were variable but low (< 0.5 μM) throughout the sampling period. Concurrent with silicate depletion, diatom abundance increased markedly, from 7.9×10^5^ cells l^-1^ at Hel_1 to 1.0×10^6^ cells l^-1^and 2.6×10^6^ cells l^-1^ at Hel_2 and Hel_3 respectively, before declining somewhat to 2.2×10^6^ cells l^-1^ at Hel_4. The diatom bloom was dominated by centric diatoms, but included a modest contribution of pennate diatoms in the latter phases of the bloom (Fig. 1a). Bacterial cell counts were relatively constant from Hel_1 to Hel_3, at 5±2 x 10^5^ cells ml^-1^, before increasing to 9.9 x 10^5^ cells ml^-1^ at Hel_4 (Fig 1b). The composition of the bacterial community, as determined using FISH staining (see below), also changed somewhat during the development of the diatom bloom, with consistently higher counts of *Bacteroidetes* (CF319a) than *Gammaproteobacteria* (GAM42a), but a stronger relative increase in *Gammaproteobacteria* from Hel_1 to Hel_4 (Fig. 1c). Counts for *Rhodobacteraceae* (ROS537) were variable but increased overall from Hel_1 to Hel_4.

**Fig. 1.**
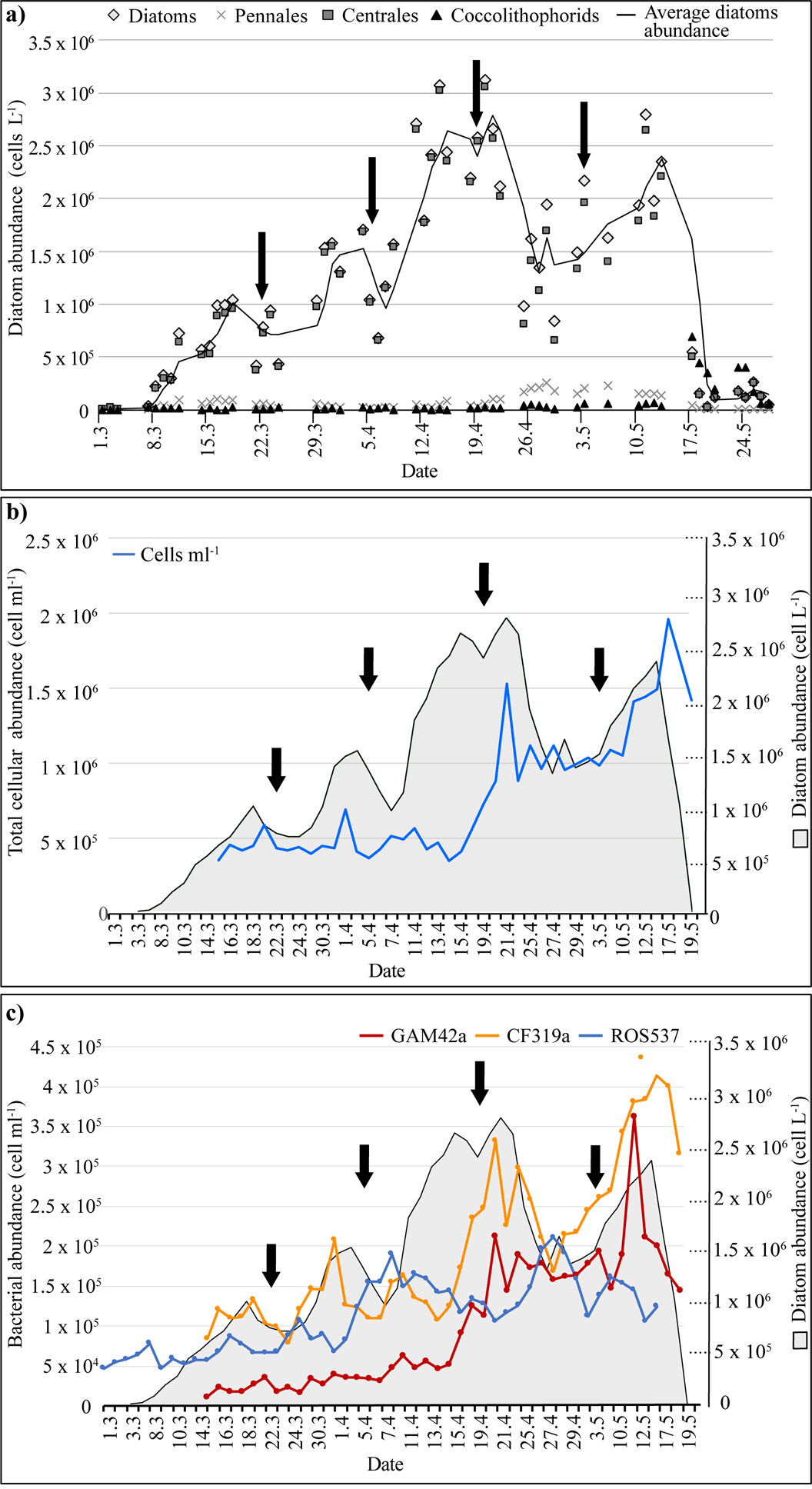
Phytoplankton and bacterial community changes at Helgoland from March through May 2016. 1a) Diatom and coccolithophorid abundance. Vertical arrows indicate the four dates (Hel_1 through Hel_4) on which samples were collected for analyses. 1b) Total cell counts (blue line) and diatom abundance (gray shaded) at Helgoland sampling site. Black arrows indicate Hel_1 to Hel_4 sampling dates. (1c): Total abundance of *Bacteroidetes* (CF319a), *Gammaproteobacteria* (GAM42a), and *Alphaproteobacteria* (ROS537) as counted by FISH. Gray shaded area shows diatom abundance, black arrows show sampling dates Hel_1 to Hel_4.

### Incubation experiments

#### Extracellular enzyme activities

We measured the hydrolysis rates and the substrate spectrum of polysaccharide hydrolase activities during the development of the bloom by adding fluorescently-labeled polysaccharides (FLA-PS) to water samples collected from Hel_1 to Hel_4 and incubating the samples over a timecourse of up to 9 days (6 days for Hel_1). The complement of polysaccharide-hydrolyzing enzymes present in heterotrophic bacteria varies widely (Zimmerman *et al.,* 2013; Xing *et al*. 2014; Kappelmann *et al.,* 2018), as does the suite of enzyme activities in natural communities in ocean waters (Arnosti *et al*., 2011; Hoarfrost *et al.,* 2019). For these incubations, hydrolysis of laminarin, xylan, chondroitin sulfate, carrageenan, and arabinogalactan was measured via changes in the molecular weight of the added polysaccharides with time as they were systematically hydrolyzed to smaller size classes (Arnosti 2003; Arnosti *et al.,* 2012).These polysaccharides were selected because they are present in the oceans, some of them in very high quantities (Alderkamp *et al*. 2007), and/or enzymes that hydrolyze them are widely distributed among marine bacteria (Alderkamp *et al*. 2007, Arnosti *et al*. 2011, Usov 2011, Xing *et al*. 2015, Kappelmann *et al.,* 2018). In addition, these polysaccharides represent a range of composition and complexity: laminarin is a glucose-containing polysaccharide, xylan is made of xylose, arabinogalactan is a mixed polysaccharide of arabinose and galactose, chondroitin sulfate is a sulfated polysaccharide of N-acetylgalactosamine and glucuronic acid, and carrageenan is a sulfated polysaccharide of galactose and 3,6-anhydrogalactose.

Enzymatic hydrolysis rates, as well as the number of substrates (the spectrum of substrates) hydrolyzed, changed notably during the development of the bloom (Fig. 2a). Initially, at Hel_1, only laminarin and xylan hydrolysis were measurable; hydrolysis rates were comparatively low (2-4 nmol monomer L^-1^ h^-1^). At Hel_2, hydrolysis rates of laminarin and xylan were somewhat higher, 6 to 8 nmol monomer L^-1^ h^-1^, rates that were maintained through the rest of the incubations. At Hel_2, chondroitin hydrolysis was first measurable at a low rate (1 nmol monomer L^-1^ h^-1^) at the 9 day timepoint. At Hel_3, carrageenan hydrolysis was first measurable at the 9 day timepoint, at a rate of 3 nmol monomer L^-1^ h^-1^ (chondroitin hydrolysis was not tested at Hel_3). By Hel_4, carrageenan and chondroitin hydrolysis were both measurable by the 6 day incubation timepoint, and the rate of chondroitin hydrolysis had increased to 6 nmol monomer L^-1^ h^-1^ by the 9 day timepoint. Thus, the spectrum of substrates hydrolyzed, the initial timepoint at which hydrolysis was detected, and hydrolysis rates of specific polysaccharides increased from Hel_1 to the later bloom stages. Intriguingly, no arabinogalactan hydrolysis was measurable at any timepoint, although cell staining indicated selfish uptake of this substrate (see below). No enzyme activities were measurable in the autoclaved seawater with added polysaccharides.

**Fig. 2a.**
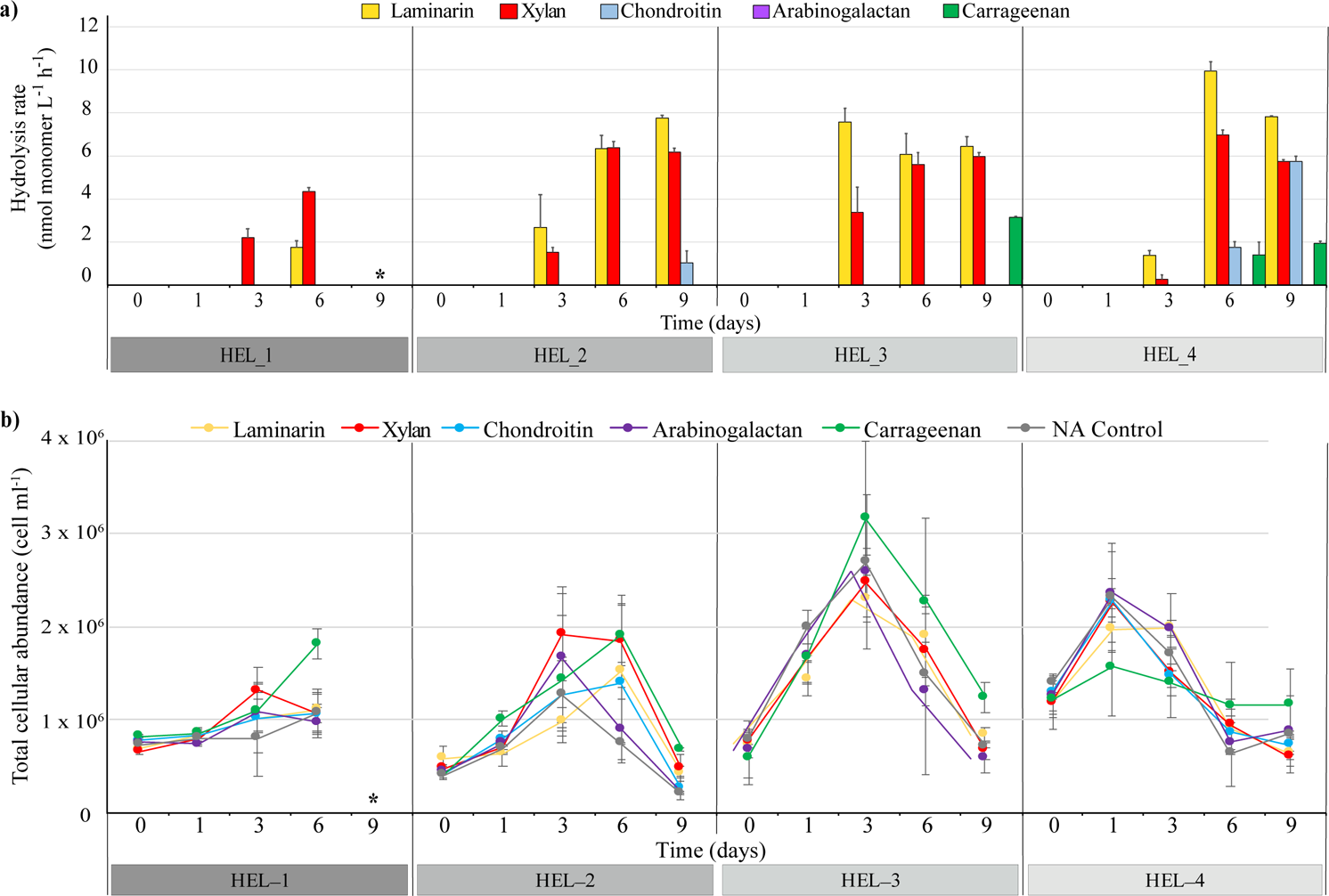
Enzymatic hydrolysis rates of laminarin, xylan, chondroitin sulfate, arabinogalactan, and carrageenan measured in triplicate incubations in each bloom phase at each sampling time point. Error bars represent the standard deviation of 3 separate incubations for each substrate. 2b) Total cell counts in the same incubations from which enzymatic hydrolysis rates were measured, plus cell counts in no addition (NA) control (no FLA-PS added). Amended incubations are color-coded (same color code as in 2a) to indicate which FLA-PS was added; error bars represent n = 6 biological and technical replicates. Note that for Hel_1, incubations lasted only 6 days. Also note that no incubations with chondroitin sulfate were carried out at Hel_3.

#### Cell counts

Cell counts in the incubation experiments varied with incubation time as well as with timepoint during the bloom (Fig. 2b). Hel_1 showed little overall change in numbers, with the exception of the carrageenan incubation, which showed a doubling of cell counts (from ca. 1 to 2 x 10^6^ cells ml^-1^). Initial cell counts for Hel_2 were lower than for Hel_1, but increased with time to the 3 or 6 day timepoint before decreasing again. Initial cell counts at Hel_3 were similar to those at Hel_1, but unlike for Hel_1, in Hel_3 cell counts in all incubations increased considerably with time. In all incubations, cell counts for Hel_3 reached a maximum of 2.5 x 10^6^ to 3.5 x10^6^ at the 3 day timepoint, and decreased to close to initial values (ca. 0.75 to 1.2 x 10^6^ cells ml^-1^) at day 9. Cell numbers for Hel_4 were initially higher (consistent with cell numbers from the environment; Fig. 1b), but peaked at day 1 for all incubations, and decreased thereafter. The temporal patterns (Fig. 2b) in cell counts were not significantly different within bloom phases for amended and unamended incubations (ANOVA: F = 0.5, p = 0.78), suggesting that overall patterns in cell numbers were driven by factors such as cell growth on natural organic matter from the bloom included in the incubations, as well as decreases in cell counts due to viruses or grazers, rather than growth in response to addition of FLA-PS to the incubations.

#### Sequencing

Against a background of changing initial microbial community composition (Fig. 1c), community composition in the FLA-PS incubations also changed with time. Next generation sequencing was carried out at selected timepoints (t0, 3d, 6d) to compare temporal changes in the incubations and to highlight possible principal responders in the incubations (Fig. 3; Figs. S2, S3). Early in the bloom (Hel_1), strong changes were seen in all incubations with time, with an especially strong response of *Bacteroidetes* (genus *Aurantivirga* and uncultured *Cryomorophyaceae*), *Gammaproteobacteria* (genus *Colwellia*), and *Alphaproteobacteria* (genera *Amylibacter* and *Planctomarina*) (Fig 3; Fig. S2). Changes observed at Hel_2 differed in part compared to Hel_1. There was again a strong response of the bacteroidetal genus *Aurantivirga*, as well as of *Ulvibacter* and uncultured *Cryomorophyaceae* (Fig. 3). The *Gammaproteobacteria* growing were affiliated with the genera *Glaciecola,* in addition to *Colwellia.* For Hel_3, a strong positive response was seen also for the alphaproteobacterial genus *Sulfitobacter*, and the gammaproteobacterial genera *Colwellia* and *Glaciecola*. At Hel_4 in particular, there was also a distinct substrate-related response, with the changes in the carrageenan and chondroitin incubations distinct from the other incubations (Fig. S3). This response included a strong increase of members of the bacteroidetal genus *Flavicella*. (Figs. 3; S2, S3). For the other substrate incubations (and for the no-addition control), the gammaproteobacterial genus *Colwellia* again increased strongly (Fig. 3, Fig. S3).

**Fig. 3:**
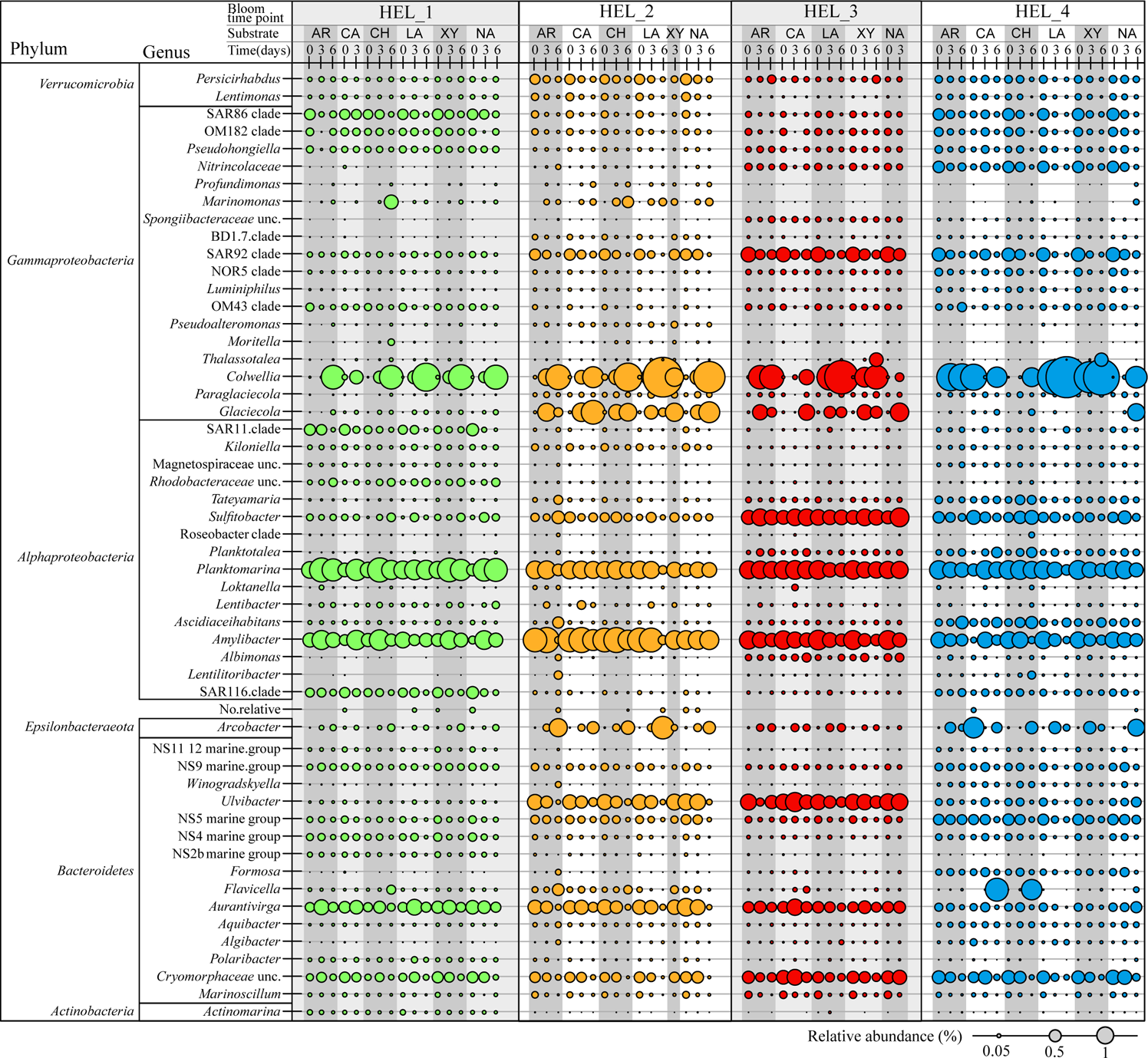
Bubble plot of bacterial genera with a minimum normalized read abundance of 0.05% at t0, 3d, and 6d for all substrates and for the unamended control incubation for Hel_1 to Hel_4. The size of the bubble represents the relative abundance (%) of each genus.

Summarizing overall community composition throughout the time course of incubation, Hel_1 (and to a lesser extent, Hel_4) showed communities that were distinctly different from the more similar Hel_2 and Hel_3 incubations (Fig. 4). Statistical analysis of the overall community compositions demonstrated that bloom phase and sampling timepoint within a bloom phase each significantly influenced overall community composition (Table 1), whereas the specific FLA-PS added (‘substrate’ in Table 1) did not. These factors - bloom phase and sampling point – had different weights for the three major phylogenetic groups considered “master recyclers” (Buchan *et al*., 2014). Among the *Bacteroidetes* and the *Rhodobacteria*, the predominant response was related to bloom phase (Hel_1 to 4; ANOSIM *R* = 0.76 and *R* = 0.56 *p*-value = 0.001, respectively). As shown by the NMDS plots in Fig. S2, all incubation timepoints of Hel_1 separated considerably from Hel_2 to Hel_4. Moreover, Hel_4 was mostly (for *Rhodobacteria*) or completely (for *Bacteroidetes*) separated from Hel_2 and Hel_3. In contrast, although the *Gammaproteobacteria* initially separated by bloom phase, over the course of the sampling timepoints they converged to the same genera (most notably *Colwellia*). Thus, sampling timepoint (rather than bloom phase) was the factor that most distinguished the *Gammaproteobacteria* (ANOSIM *R =* 0.55, *p-*value = 0.001, Fig. S2).

**Fig. 4:**
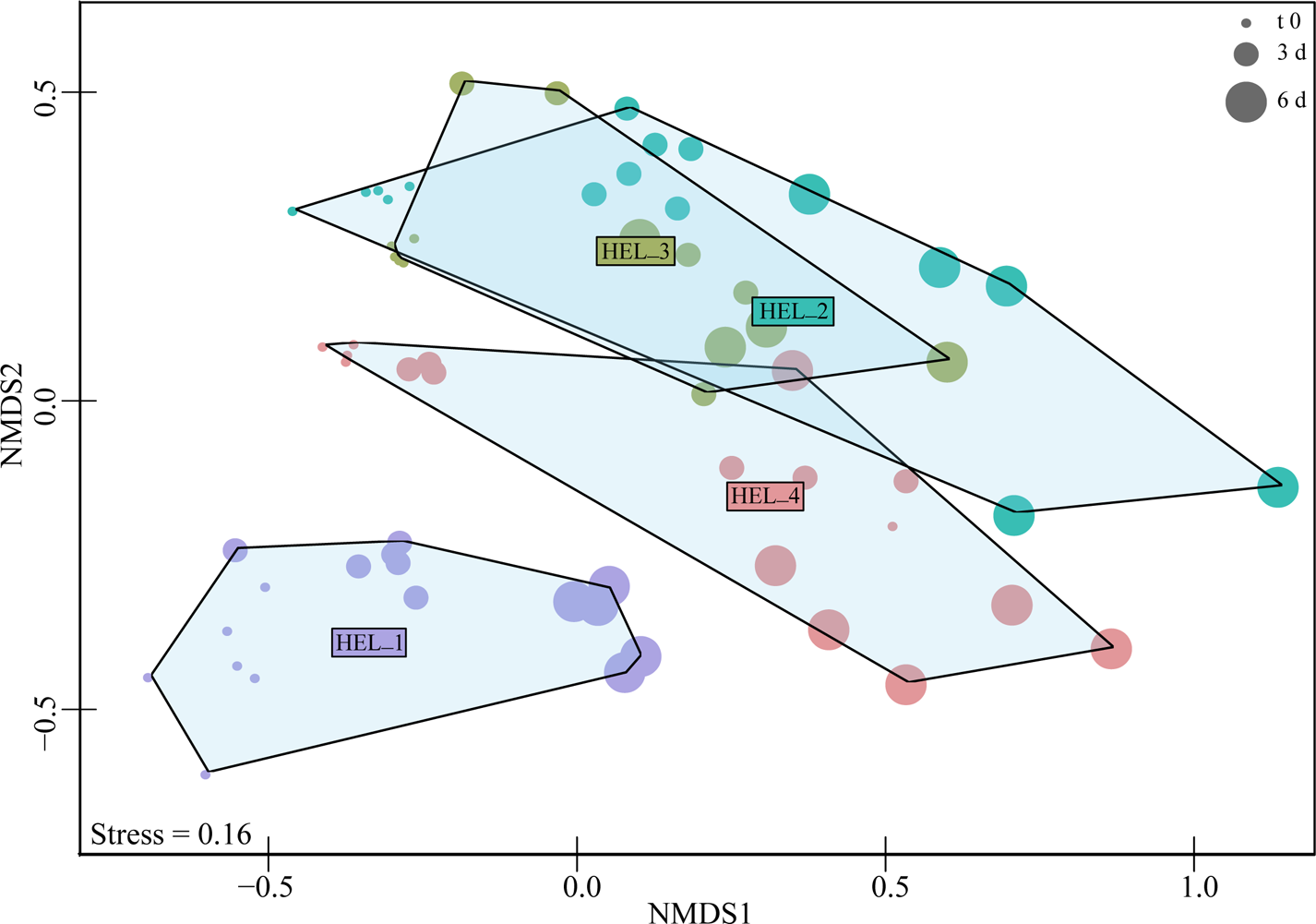
NMDS ordination based on Bray-Curtis dissimilarity for the comparison of microbial community composition between 4 bloom phases (color coded). Hel_1 separates strongly from the other timepoints, and Hel_4 is mostly separated from Hel_2 and Hel_3. The time course of incubation for each sample time are indicated by increasing bubble size.

**Table 1.** Permutational multivariant analysis of variance (PERMANOVA) and analysis of similarly (ANOSIM) of bacterial community composition based on Bray–Curtis dissimilarities of relative read abundance, showing significant differences among bloom phases, and across individual time points within a bloom phase. There were no significant differences by substrate.

| Source of variance | PERMANOVA |  |  |  | ANOSIM |
| --- | --- | --- | --- | --- | --- |
|  | d.f | SS | pseudo F | R<sup>2</sup> | R |
| Substrate | 5 | 0.39 | 11.03 | 0.05 | -0.03361 |
| Bloom phase | 1 | 17.62 | 248.8 | 0.23* | 0.3508* |
| Sampling timepoint | 1 | 13.34 | 188.67 | 0.18* | 0.5812* |
| Residuals | 57 | 40.36 |  | 0.54 |  |
| Total | 64 | 75.24 |  | 1 |  |

PERMANOVA test show influence of factors: "substrate", "bloom phase", and "sampling timepoint" on community composition, *p*-values were obtained using sum of squares and 999 permutations. ANOSIM performed with 999 permutation. d.f. degrees of freedom, SS sum of squares, \* denotes significance ( $p < 0.001$ ).

#### Bacterial staining by FLA-PS and FISH

The abundance and activity of ‘selfish’ bacteria was determined by tracking cellular uptake of FLA-PS in the same incubations set up for enzyme activity measurements (Reintjes *et al*., 2017). Selfish uptake was substrate as well as bloom-phase dependent. At all bloom phases, laminarin was consistently taken up by a larger fraction of the bacterial community than the other substrates (Fig. 5). The fraction of cells taking up laminarin increased from 14% at Hel_1 to a maximum of 29% at Hel_2. It remained high with maxima of 20% and 25% of total DAPI-stained cells at Hel_3 and Hel_4, respectively. The fraction of cells taking up xylan, chondroitin, and arabinogalactan also was higher at timepoints after Hel_1, but the fraction of total cells taking up a substrate was considerably smaller than for laminarin: for xylan, maximum uptake was 14% for at Hel_3, for chondroitin the maximum was slightly over 5% for Hel_2 (no incubations with chondroitin for Hel_3), and maximum arabinogalactan uptake was 12% for Hel_2 (Fig. 5). Note that carrageenan uptake could not be counted, since the background fluorescence from this gel-forming substrate was too high. Overall, selfish staining was highest at Hel_2 (Fig. 5).

**Fig. 5:**
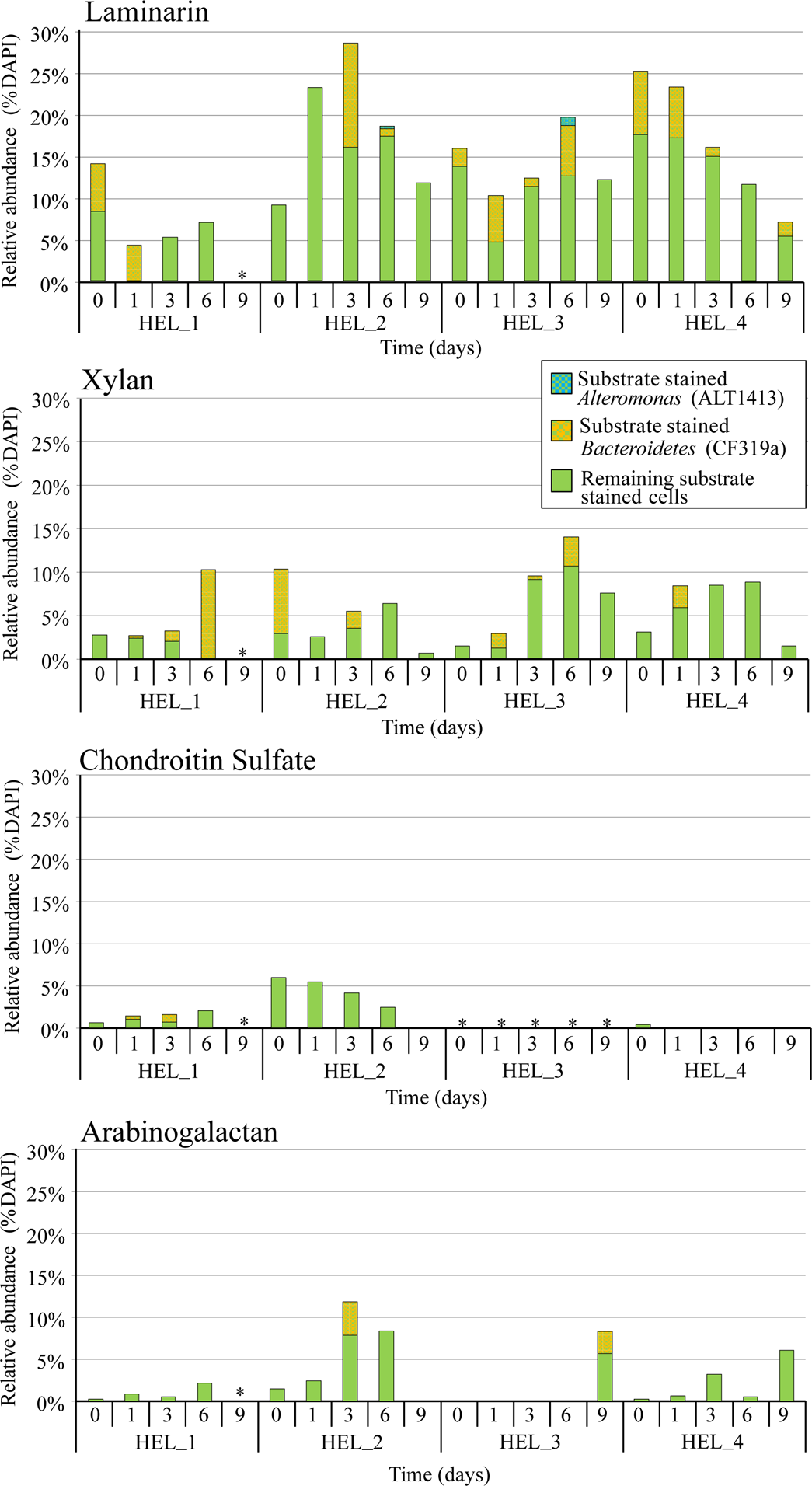
Total cellular abundance of FLA-PS stained cells (selfish uptake) for all incubations and timepoints. A fraction of these selfish cells were identified using FISH as members of *Alteromonas* (blue; ALT1413) and *Bacteroidetes* (yellow; CF319a); the remaining substrate stained cells are indicated by green color.

FISH staining was carried out using the probes CF319a and ALT1413 to identify the overall presence of *Bacteroidetes* and *Alteromonadales* in the incubations. In the laminarin incubations, a high fraction (10 to 40% of DAPI-stainable cells) were CF319a positive (Fig. S4). These incubations also had a typically smaller and more variable fraction of ALT1413 positive cells, ranging from nearly zero for Hel_1 to a high above 30% for Hel_2 to 3%-20% for Hel_3 and Hel_4 (Fig. S4). This pattern of variable proportions of ALT1314-stainable cells and a generally high fraction of CF319a-positive cells also characterized the xylan, chondroitin, and arabinogalactan incubations. Carrageenan incubations were not analyzed using FISH due to high background fluorescence.

A variable proportion of these ALT1413 and CF319a-stained cells were identified as ‘selfish’ (Fig. 5). The fraction of FLA-PS-stained (‘selfish’) cells that were not CF319a/ALT1413 positive was frequently one-third to one-half of the total, thus there are additional clades with a selfish polysaccharide uptake mechanism (Fig. 5). The microscopic evidence of ‘selfish’ uptake of arabinogalactan (Fig. 5) was particularly intriguing, since there was no evidence of external hydrolysis of this substrate (Fig. 2a).

## Discussion

Spring phytoplankton blooms in temperate environments fuel the base of complex food webs, producing a rapid increase in organic matter that is recycled in the upper ocean as well as exported below the thermocline (Boss and Behrenfeld, 2010). The progression of phytoplankton during the course of the bloom is mirrored by a succession of bacterial community members whose dynamics are also complex (Riemann *et al*., 2000; Teeling *et al*., 2012; 2016). These successional patterns are likely the result of finely-tuned bacterial responses to changing organic matter availability derived from the phytoplankton. Specialist heterotrophic bacterial groups vary in their genomic repertoire (Bauer *et al*., 2006; Xing *et al*., 2014; Badur *et al*., 2017; Unfried *et al*., 2018; Kappelmann *et al*., 2018; Krüger *et al*., 2019), and thus in their ability to access complex substrates produced in abundance by phytoplankton; the progression during a bloom of autotrophic and heterotrophic members of the plankton is therefore closely intertwined (Buchan *et al.,* 2014). Measuring two distinct mechanisms of substrate processing while tracking dynamic changes in microbial community composition provides new insight into the conditions under which different strategies of substrate processing prevail. To arrive at a clear picture of these intertwined dynamics, however, first the changes in community composition, and then the changes in substrate processing mechanisms need to be examined.

### Community compositional changes

Our incubation experiments were in effect small bottle experiments, in which a diverse array of microorganisms present in the water sampled grew primarily in response to the natural organic matter present (Fig. 2b; Figs. S2, S3). As demonstrated by the similarity in cell counts between the amended and unamended incubations, overall changes in bacterial cell numbers during our incubations (Fig. 2b) were more closely related to the four bloom phases and the bloom-derived organic matter naturally included in the incubations, and not to the addition of a comparatively low concentration of FLA-PS to a given incubation. In terms of bacterial community composition, as expected (Buchan *et al*., 2014), members of the *Alphaproteobacteria*, *Gammaproteobacteria*, and *Bacteroidetes* were among the most prominent responders (Fig. 3, Figs. S2, S3). The nature of the responses among these groups varied, however. The development of dominant *Bacteroidetes* and *Alphaproteobacteria* genera separated clearly among bloom phases, likely as a result of changes of bacterial community composition over the course of the bloom (Fig. 3, Fig. S2). Although the *Gammaproteobacteria* present at t0 (Fig. S2) also differed by bloom phase, the specific increase in *Colwellia* in all bloom phase incubations decreased the overall gammaproteobacterial community dissimilarity, and thus drove the overall composition from different initial starting points in a similar direction during the course of the incubations (Fig. S2). This strong response is not surprising, as *Colwellia* is regarded as a ‘boom and bust’ specialist (Teira *et al.,* 2008), highly responsive to the presence of complex organic matter, including in amendment incubations with diatom-derived high molecular weight organic matter in water from the North Atlantic (Balmonte *et al*., 2019), and in response to of diatom-derived dissolved organic matter in continuous cultures experiments in the Southern Ocean (Landa *et al*., 2018). Other organisms that have been reported to respond to diatom-bloom-related DOM include *Alphaproteobacteria* of the genus *Sulfitobacter*, and the genus *Ulvibacter* of the *Bacteroidetes* (Landa *et al*., 2018), which were present in all of our incubations, including the no-addition control (Fig. S2).

### Bacterial functional responses to bloom dynamics

In addition to the compositional response, the development of the bloom triggered a clear functional response among the bacterial community, characterized by an increase in potential polysaccharide hydrolysis rates and a broadened spectrum of polysaccharide hydrolase activities between the early and the later phases of the bloom (Fig. 2a). These responses are likely due to increased quantity as well as higher complexity of organic matter available during the course of the bloom (Krüger *et al*., 2019). Hel_1 was taken early in the first diatom bloom after the winter, conditions under which little bioavailable HMW organic matter would be present. We suggest that over the course of the next 6 weeks, the concentrations and the complexity of the polysaccharides increased, simultaneous with the complexity of the phytoplankton community as the Centrales-type diatoms dominant in Hel_1 transitioned to a mixture containing Pennales and coccolithophorids that was still dominated by Centrales (Fig. 1a). As the bloom progressed, the glycan pool diversified, containing increasing quantities of polysaccharides of higher complexity as well as freshly released polysaccharides. The early bloom phase showed hydrolysis only of laminarin and xylan, which are compositionally less complex than chondroitin and carrageenan; the hydrolysis of these latter two substrates began only after the initial phase of the bloom (Fig. 2a). These developing responses to the presence of specific polysaccharides likely reflected transitions in the bloom-associated microbial community, with bacteria possessing the highly specific enzymes required to depolymerize more complex substrates increasingly present as the bloom progressed (Teeling *et al*., 2016; Krüger *et al.,* 2019). Thus, in order to observe hydrolysis of specific complex substrates in our incubations, a starting community for a given incubation had to contain sufficient member(s) with the capability of responding to that complex substrate. Our measurement of activities from a wider spectrum of enzymes with progressing bloom stage is consistent with previous data from Helgoland spring blooms showing increased expression of proteins associated with polysaccharide degradation with time (Teeling *et al*., 2012).

The incubations with carrageenan and chondroitin highlight the intertwined roles that bacterial community capabilities and substrate complexity play in determining the fate of bloom-derived organic matter. Both carrageenan and chondroitin were hydrolyzed only after the initial bloom phase; sequencing of the carrageenan and chondroitin incubations in the latter phase of the bloom revealed communities that were distinctly different – especially among members of the *Bacteroidetes* (Fig. 3, Figs. S2, S3) - from those in the laminarin, xylan, and control incubations at the same timepoint, as well as from incubations with carrageenan and chondroitin early in the bloom (Fig. S3). These two substrates provide an example of a substrate-driven effect on microbial community composition (Landa *et al*., 2013). The prominent response later in the bloom of the bacteroidetal genus *Flavicella* (Fig. 3, Fig. S3) to both carrageenan and chondroitin, moreover, suggests that it may have a role in the degradation of these complex, charged substrates, in accordance with the idea that certain heterotrophs are selectively able to utilize organic matter from specific phytoplankton (Sarmento *et al*., 2012).

The heterotrophic bacterial community responded to the development of the bloom not only with increased activities of extracellular enzymes that yield lower molecular weight polysaccharide fragments outside the cell; some members of the community also increased ‘selfish’ uptake of substrates. Laminarin uptake, which is readily induced (Reintjes *et al*., 2017), was high already for Hel_1, with 14% of DAPI-stained cells binding this FLA-PS at t0 (∼ 15 minutes after substrate addition; Fig. S5). A considerable fraction of bacteria incorporated laminarin at t0 at subsequent bloom sampling times, reaching a maximum slightly over 25% of total cell counts at t0 for Hel_4. Through the course of each bloom incubation, the fraction of cells taking up laminarin varied by time point (Fig. 5), but overall remained at a high level. Uptake of chondroitin and arabinogalactan changed considerably during the bloom, with initial values less than 3% of total cell counts throughout the Hel_1 incubation, increasing to ca. 6-12% at Hel_2, and decreasing to somewhat lower values (arabinogalactan) to very low levels (chondroitin) for later bloom phases.

Xylan uptake changed less between bloom sampling times, typically reaching a maximum of 8-14% of total cells at some point in each of the incubation series, although initial uptake (at all but Hel_2) was low. The percentage of total cells taking up laminarin and xylan is similar to previous measurements in the northern temperate province of the North Atlantic (Reintjes *et al*., 2017); chondroitin and arabinogalactan uptake overall are somewhat lower than previously seen in the northern temperate province (Reintjes *et al*., 2019).

Consideration of the ‘selfish’ bacteria that became stained with FLA-PS reveals several intriguing features. No hydrolysis of arabinogalactan was measured, yet 1-12% of total cells showed ‘selfish’ uptake of this substrate, with overall higher uptake percentages after Hel_1 (Fig. 5). In these incubations, therefore, arabinogalactan processing appears to have occurred solely via selfish pathways, with no detectable production of low molecular weight hydrolysis products in the external environment. This pattern of solely-selfish uptake contrasts with our previous investigation, in which both external hydrolysis and selfish uptake of arabinogalactan was measurable at all five stations in the North and South Atlantic (Reintjes *et al*., 2019). In that investigation, however, fucoidan was also taken up in a ‘selfish’ manner, with no detectable extracellular hydrolysis (Reintjes et al., 2019). Possibly there are links between selfish uptake and initiation of extracellular hydrolysis, particularly for the more complex substrates. If selfish uptake is also coupled to low production of external hydrolysate (as has been shown for some members of the *Bacteroidetes*; Rakoff-Nahoum *et al*., 2016), then perhaps a threshold concentration of arabinogalactan hydrolysis products was not reached to initiate such activity at Helgoland.

Another intriguing observation relates to our efforts to identify the selfish bacteria by FISH. The term selfish uptake has been coined for gut *Bacteroidetes* (Cuskin *et al*., 2015); again in this study, a fraction of the marine *Bacteroidetes* – mostly members of the class *Flavobacteriia* - were shown to behave selfishly with respect to the substrates tested (Fig. 5). Not surprisingly, even comparatively structurally simple glycans such as laminarin are not taken up by all marine *Bacteroidetes* (Fig. S4); comparative genome analysis of 53 strains of marine *Bacteroidetes* suggested that only 33 have the canonical polysaccharide utilization loci for laminarin (Kappelmann *et al*., 2018). The same consideration applies to *Alteromonas,* where a very small fraction of the cells was stained by FLA-PS (Fig. 5). In both cases, part of the underlying cause for unstained cells may be that the respective genes are absent or not induced; insufficient FISH probe coverage is also a possibility. At most timepoints and for most incubations, a sizable fraction ranging from 45% to 100% of the selfish bacteria were not detected with the FISH probes used. This observation suggests that bacteria other than *Bacteroidetes* and a few members of the *Gammaproteobacteria* may be involved in the uptake of FLA-PS. This point has already been demonstrated for *Verrucomicrobia* (Martinez-Garcia *et al*., 2012) and *Planctomycetes* (Reintjes et al., 2017). These other ‘selfish’ bacteria must transport polysaccharides using mechanisms other than the well-studied Sus systems (starch utilization system; Cho and Salyers, 2001).

### Interacting strategies of carbon cycling during a spring bloom

Multiple organisms and factors interact during a phytoplankton bloom, changing the quantity and nature of organic matter produced and consumed by heterotrophic microbial communities, and also the strategies used to consume fractions of differing complexity. Here, it is important to note that measurements of selfish bacteria sampled immediately (∼ 15 minutes) after substrate addition likely reflect the conditions in the field during the bloom. Looking at these time-zero points, a clear distinction between laminarin on the one hand and the other substrates on the other is evident (Fig. S5). Initial laminarin uptake was high at Hel_1 (14% of total cell counts), and became higher during the course of the bloom, reaching 25% at Hel_4. For arabinogalactan, chondroitin, and xylan, initial uptake at time zero at Hel_1 was low (ranging from 0 to 3%), increased substantially at Hel_2, and decreased again at Hel_3 and Hel_4 to 0 to 3% of total cells. Integration of these data in a schematic figure (Fig. 6a) shows that the early bloom phase (Hel_1) is characterized by low selfish uptake, except for laminarin, and a limited range of polysaccharide hydrolase activities (Fig. 2a). Two weeks further into the increasing bloom (Hel_2), more – and most likely more complex – organic matter is available as diatom numbers rapidly increase, and are turned over by grazers and viruses (Fig. 1a). Selfish organisms are now more numerous and diverse with respect to the substrate transported. They presumably have had sufficient exposure to a broader range of substrates such that selfish uptake is induced or that selfish bacteria have increased in relative abundance (Reintjes *et al*., 2017). Extracellular hydrolysis rates are also higher, and a slightly wider spectrum of substrates is hydrolyzed. At the late bloom stage (Hel_4), even more organic matter of greater complexity is available, and bacterial cell numbers have increased substantially (Fig. 1b). This increase must have included bacteria that carry out external hydrolysis, since a broad spectrum of substrates is hydrolyzed. Selfish uptake remains high primarily for laminarin, but much less so for the other substrates, presumably because the selfish bacteria doubled less rapidly than the external hydrolyzers in response to the rapid increase in organic matter, or they were killed by viruses or grazers. At this stage, external hydrolyzers become more important for the degradation of the more complex substrates.

**Fig. 6a:**
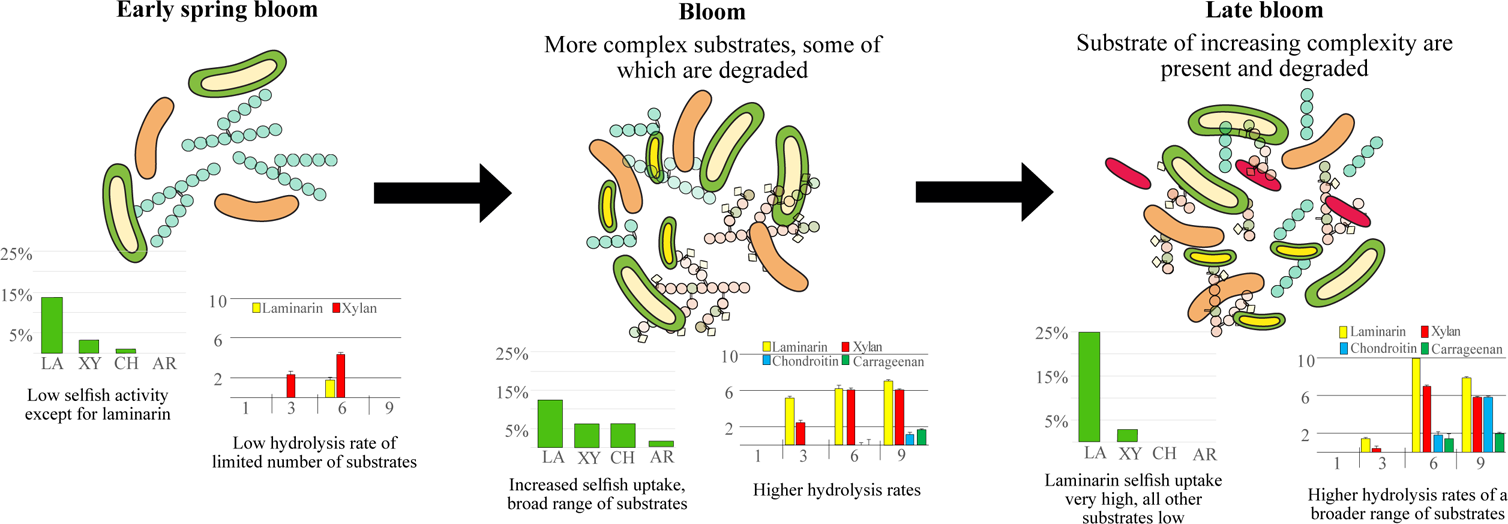
*The relationships between substrate complexity and quantity, selfish uptake, and extracellular hydrolysis changes substantially during the course of a bloom*, as demonstrated by external hydrolysis rates (same data as in Fig. 2a) and selfish uptake at t0 (plots from Fig. S5), shown below the main schematic diagram. During the early bloom phase, both external hydrolysis and selfish uptake are low, except for laminarin, a widely-available substrate. During the bloom phase, more complex substrates also become available; a broader range of substrates are taken up via a selfish mechanism; external hydrolysis rates increase somewhat. The late bloom phase is characterized by greater quantities of more complex substrates that support bacteria carrying out external hydrolysis; these bacteria double quickly. Fewer selfish bacteria are supported, since much of the substrate is hydrolyzed externally to low molecular weights. Because of the high quantity of laminarin available throughout the bloom, it is processed by both selfish and external hydrolysis even in the late bloom phase (see Fig. 6b).

Short-term changes in polysaccharide utilization mechanisms are likely driven in part by the quantity and complexity of available substrates: initial exposure to readily-dissolved substrates such as laminarin can lead to rapid selfish uptake, and hoarding of substrate for those cells. When organic matter supply is sufficient to fuel a larger community and is enriched in particulate organic matter, external hydrolysis becomes more prevalent (Fig. 6b). It is intriguing that laminarin is taken up by a selfish mechanism throughout the bloom, and that the fraction of selfish uptake increases even as external hydrolysis increases. This situation likely reflects the fact that in diatom-dominated blooms, laminarin is constantly released in large quantities. Global production of laminarin is estimated to be on the order of 5-15 Gt annually (Alderkamp *et al*., 2007); it therefore must constitute a major energy source for many marine bacteria.

**Fig. 6b:**
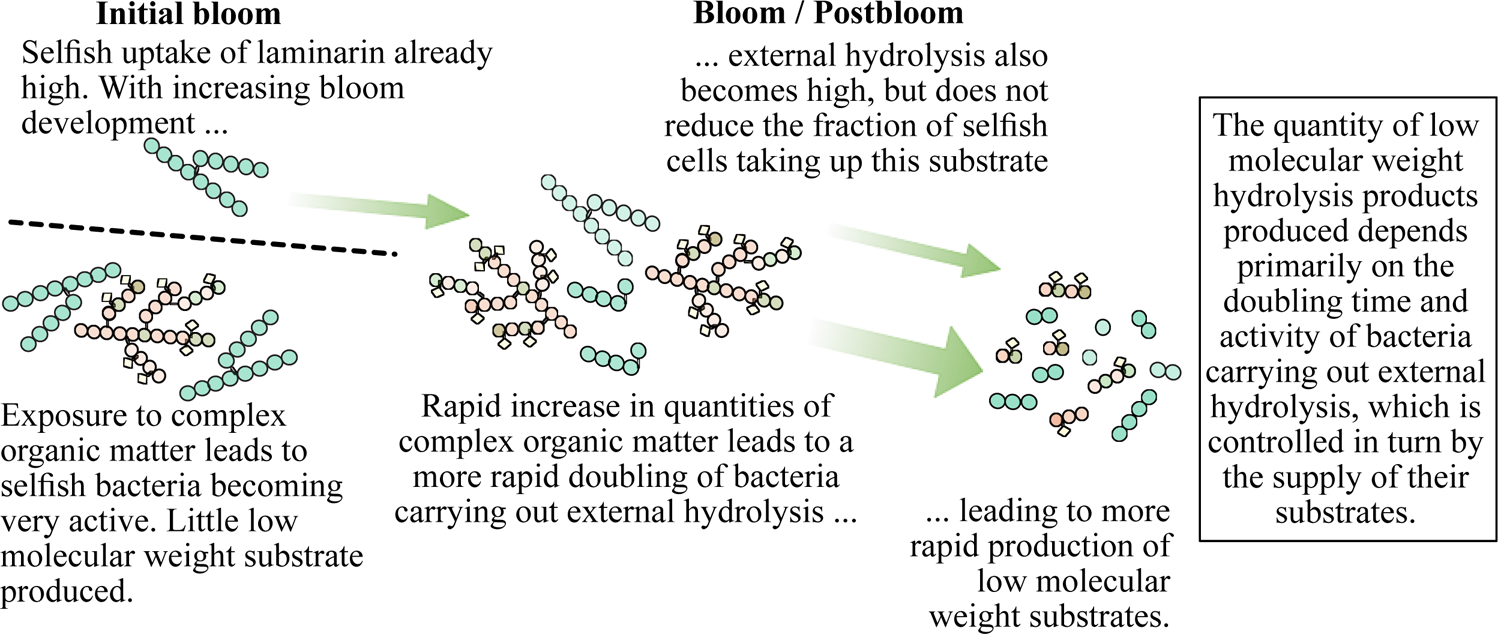
*Substrate complexity and quantity both affect the nature of substrate processing.* In the early bloom stages, selfish bacteria become more active as a wider range of substrates are initially made available. Increasing quantity of these substrates, however, leads to more rapid doubling of bacteria that carry out external hydrolysis; in late bloom stages, external hydrolysis is the more dominant substrate processing pathway. Because of the high quantity of laminarin available throughout the bloom, however, it supports both selfish uptake and external hydrolysis in all bloom phases. The net outcome for organic matter processing is that overall more low molecular weight substrates are available in the external environment in the bloom/postbloom stages.

These interactions between selfish and sharing bacteria have major implications for downstream carbon flow: scavenging bacteria profit from low molecular weight substrates provided by the external hydrolyzers, but not - for the most part - from the activities of selfish bacteria (Arnosti *et al*., 2018). The quantity and type of substrates available to feed the downstream bacterial foodchain thus are closely linked to these upstream processes. Sarmento and Gasol (2012) found that uptake of dissolved organic carbon (DOC) from a variety of phytoplankton varied markedly among major phylogenetic groups of heterotrophic bacteria, and that with an increase in DOC quantity, quality became less important (Sarmento *et al*., 2016). We find that the mode of substrate processing – selfish or external hydrolysis – varies markedly by substrate as well as by bloom phase, i.e. by substrate quality, by substrate quantity, and also by the bloom-stage dependent composition of the heterotrophic bacterial community. Quality determines the specific enzymatic requirements for selfish bacteria, as well as abundance of ‘specialist’ bacteria among external hydrolyzers, as seen for carrageenan and chondroitin. Quantity seems to be important in fueling the growth of external hydrolyzers that produce a broader range of hydrolysis products that presumably fuel scavenging bacteria as well as the external hydrolyzers: with higher quantities of a substrate, a wider fraction of the total heterotrophic community can benefit. The bloom-stage composition of the heterotrophic bacterial community helps determine the predominance of the individual mechanisms of substrate utilization, based on genomic specialization and linked community-level interactions.

### Experimental procedure

#### Sampling and Substrate Incubations

Seawater samples were collected near the island of Helgoland in the German Bight at the long-term monitoring station Kabeltonne (54°11’03” N, 7°54’00” E). 25 L of seawater was collected between 10:00 - 12:00 on March 22^nd^, April 5^th^, April 19^th^, and May 3^rd^, 2016. The samples are referred to as Hel_1 to Hel_4, respectively. Due to the abundance of organic matter, samples Hel_2, Hel_3 and Hel_4 were prefiltered through a 142 mm diameter 10 µm pore-size polycarbonate filter (Milipore) by peristaltic pump (<200 mbar) for the incubations (see below).

At each sampling date, fifteen 650 ml subsamples were placed into acid-washed sterile glass bottles and incubated with one of five fluorescently labelled polysaccharides (FLA-PS) for a total of 9 days, with the exception of Hel_1, for which the incubations only lasted for 6 days. Single substrates (fluoresceinamine-labelled laminarin, xylan, chondroitin sulfate, arabinogalactan, and carrageenan) were added at a concentration of 3.5 µM monomer equivalent. Additionally, a no-addition control, consisting of a 650 ml subsample incubated in a sterile glass bottle without added polysaccharide, and five killed controls, consisting of 50 ml subsamples of autoclaved seawater incubated with polysaccharides, were taken at each time point (0, 1, 3, 6, and 9 days). All of the enzyme activities were monitored for Hel_1, Hel_2, and Hel_4; due to a lack of available substrate, chondroitin sulfate was not included in the measurements for Hel_3.

All bottles (15 incubations, 1 no-addition incubation, and 5 killed controls per sampling time point) were incubated at *in situ* temperatures (6°C, 7°C, 8°C, 8.5°C, for Hel_1 to Hel_4, respectively) in the dark. At regular intervals subsamples were collected from the incubations (typically ca. 15 min (referred to as t0), 1d, 3d, 6d, and 9d). For microscopy and FISH, 20 ml of water was fixed with 1% formaldehyde for 1 h at room temperature and subsequently filtered through a 47 mm (0.2 µm pore size) polycarbonate filter, applying a gentle vacuum of < 200 mbar. After drying, the filters were stored at −20°C until further analysis. For microbial diversity analyses, 10 ml of water was filtered through a 25 mm (0.2 µm pore size) polycarbonate filter using a Whatman 420200 Swin-Lok filter holder (Sigma-Aldrich, Munich, Germany) and stored at −80°C until further analysis. Two ml of the filtrate from the microbial diversity sample was collected and stored at −80°C for measurement of extracellular enzyme activities.

#### Physical/chemical data and phytoplankton abundance

Phytoplankton abundance and composition, as well as physical and chemical data (salinity, temperature, chlorophyll *a*, silicate, nitrate/nitrite, phosphate) were measured as part of the Helgoland Roads LTER time series, as described in detail in Teeling *et al*. (2012). These time series data are available as part of the open database Pangaea (http://www.pangaea.de) https://doi.pangaea.de/10.1594/PANGAEA.864676.

#### Preparation and characterization of FLA-PS and measurement of extracellular enzyme activities

Fluorescently labeled polysaccharides (FLA-PS) laminarin, xylan, chondroitin sulfate, arabinogalactan, and lamda-carrageenan were synthesized and characterized following the procedure described in Arnosti (2003). In brief, polysaccharides (Sigma-Aldrich) were dissolved in milli-Q water, activated with CNBr, injected onto a column of Sephadex gel (G-25 or G-50, depending on the molecular weight of the polysaccharide), and the initial peak (followed via absorbance at 290 nm using a UV-Vis detector) was collected into a vial containing fluoresceinamine. The labeled polysaccharide was separated from unreacted fluorescent tag via an additional round of column chromatography or by using Vivaspin 20 concentrators (Vivaproducts). Arabinogalactan was labeled using the procedure described (Arnosti 2003) for chondroitin sulfate and laminarin. Lamda-carrageenan was also labeled using this procedure, but only 10 mg of lamda-carrageenan were initially dissolved in 2 ml milli-Q water, and the solution was filtered through a 0.8 um pore-size disposable filter that was rinsed twice with 500 ul milli-Q water to constitute the initial polysaccharide solution for activation. FLA-PS were chemically characterized by measuring carbohydrate content (Chaplin and Kennedy, 1986) and fluorescent tag abundance via absorbance at 490 nm.

Extracellular enzymes are measured by following the changes in substrate molecular weight with incubation time (Arnosti, 2003). Since the extent of enzyme activity cannot be determined *a priori*, samples are collected over a time course and analyzed. As described in considerable detail (and shown in chromatograms in Arnosti (2003)), hydrolysis of a FLA-PS is detected as the change in substrate molecular weight as the polysaccharide is hydrolyzed to progressively smaller sizes with time; initial hydrolysis of a polysaccharide can be detected only when a sufficient fraction of the added polysaccharide pool has shifted in molecular weight class. For this reason, there is a lag between initial occurrence and initial detection of hydrolytic activity. Samples from the time course incubations (samples collected at 0, 1, 3, 6 and 9 days; for Hel_1, incubations were not run beyond 6 days) were injected onto a system consisting of two columns of Sephadex gels connected in series with column effluent passing through a Hitachi fluorescence detector set to excitation and emission wavelengths of 490 and 530 nm, respectively. Columns are standardized with commercially-available FITC dextran standards (50 kDa; 10 kDa; 4 kDa; monomer; free tag; all from Sigma) so the elution time corresponding to different molecular weight classes is known. The extent to which hydrolysis can be detected is limited by the resolution of the gel permeation chromatography system, since hydrolytic activity is only detectable when a measurable fraction of the added polysaccharide pool is shifted to a lower molecular weight class.

#### Substrate Staining, FISH and Automated Microscopy

All cell staining, FISH, and microscopy was done as described in detail in Reintjes *et al*. (2017). Briefly, total cell counts were determined by staining with 4’,6-diamidino-2-phenylindole (DAPI). Cells were counted using the automated image acquisition and cell couting system described by Bennke *et al*. (2016), and verified by manual counting. The number of substrate stained cells were determined by enumerating cells which had both a positive DAPI signal and a positive substrate signal (excitation 488 nm wavelength, minimum signal background ratio of 2.5 at constant exposure time of 200 ms). Subsequently, two group specific FISH probes targeting the *Bacteroidetes* (CF319a (5’-TGG TCC GTG TCT CAG TAC −3, formamide 35%, accession no. pB-42, (Manz *et al.,* 1996), and *Alteromonadaceae* (ALT1413 (5’-TTT GCA TCC CAC TCC CAT −3’, formamide 40%, Accession no. pB-609; Pernthaler *et al*., 2002)) were applied to quantify the abundance of these groups during the incubations. All cell counts and microbial abundance data are openly available at https://doi.pangaea.de/10.1594/PANGAEA.903579.

#### DNA Extraction, Polymerase Chain Reaction (PCR), Size Selection and Quantification

DNA extractions were done using the MoBio Power Water DNA Extraction Kit (MoBio Laboratories, Inc., CA, USA) as recommended by the manufacturer. PCR was carried out using the Platinum PCR SuperMix High Fidelity polymerase kit (Thermo Fisher), using the primers S-D-Bact-0341-b-S-17 and S-D-Bact-0785-a-A-21 targeting the V3-V4 variable region of the 16S rRNA, evaluated by Klindworth *et al.,* (2013). Both primers were fusion primers with additional adaptor and barcode sequences at the 5’ end to allow sequencing and separation of samples in down-stream analyses. The reverse primers contained the Ion tr-P1 adaptor and the forward primers contained both the Ion A adaptor and one of 40 IonXpress barcodes (Ion Xpress 1 - 40) as well as the key sequence (GAT) before the primer. Reverse fusion primer sequence: (5’-CCT CTC TAT GGG CAG TCG GTG AT GAC TAC HVG GGT ATC TAA TCC −3’). Forward fusion primer sequence: (5’-CCA TCT CAT CCC TGC GTG TCT CCG ACT CAG - barcode sequence - GAT CCT ACG GGN GGC WGC AG-3’). The primers were barcoded with the Ion Express^TM^ Barcodes (ThermoFisher) number 1-40.

After amplification the PCR products were size selected using Agencourt AMPure XP (BeckmanCoulter, Krefeld, Germany). All template libraries and final sequencing pools were analysed on a fragment analyser (AATI, USA) using the DNF - 472 standard sensitivity NGS kit sizing DNA (AATI, size range from 25 bp – 5,000 bp and up to a minimum of 0.1 ng µl^-1^) as recommended by the manufacturer

#### Ion Torrent Sequencing and Raw Sequence Processing

All substrate incubations of Hel_1, to Hel_4 for time points t0, 3 days, and 6 days were sequenced using an Ion Torrent PGM platform (Thermos Fisher). Library preparation was done as recommended by the manufacturer using an Ion OneTouch 2 Instrument (Thermo Fisher), Ion OneTouch ES instrument (Thermo Fisher) and the Ion PGM Hi-Q View OT2 kit. Subsequently, the libraries were sequenced on an Ion Torrent PGM sequencer (Thermo Fisher) using the Ion PGM Hi-Q View Sequencing Kit (Thermo Fisher) and Ion 314 chip v2 (Thermo Fisher) with a total of 1200 flows per sequencing run.

The raw reads were quality trimming using user define “stringent” settings in the Torrent Suite software (Thermo Fisher, Stringent settings:--barcode-mode 1 --barcode-cutoff 0 --trim-qual-cutoff 15 --trim-qual-window-size 10 --trim-min-read-len 250). The remaining high quality reads were classified using the standard settings of the SilvaNGS pipeline (Quast *et al*., 2013). Briefly, the pipeline processing involved alignment against the SSU rRNA seed of the SILVA database release 132 using SINA v1.2.10. (Quast *et al*., 2013) and subsequent quality controls for sequence length (> 200 bp), minimum quality score (30), minimum alignment score (40), minimum alignment identify (40%), maximum ambiguities (< 2%) and maximum homopolymers (< 2%). The remaining reads were then de-replicated and clustered using cd-hit-est (version3.1.2; Li and Godzik, 2006), running in accurate mode, ignoring overhangs and applying identity criteria of 1.00 and 0.98, respectively. Then the classification is performed by a local nucleotide BLAST search against SILVA SSURef 132.1 NR database using blast −2.2.22+ with standard settings. A detailed description of the SILVAngs project and pipeline can be found (https://w.arb-silva.de/ngs/Index.html#about: Quast *et al.,* 2013).

All sequence data was deposited in the European Nucleotide Archive (ENA; Toribio *et al.,* 2017) using the data brokerage service of the German Federation for Biological Data (GFBio, (Diepenbroek *et al*. 2014), in compliance with the MIxS standard (Yilmaz *et al*. 2011). The INSDC accession number for the data is PRJEB33503.

### Statistical Analysis

All statistical analyses and graphing of the microbial diversity data were done using RStudio version 1.2.335 with the packages Picante version 1.8, Rioja version 0.9-21 and RVAideMemoire version 0.9-73 (RStudio Team, 2018; Juggins, 2016; Kembel *et al*., 2019; Herve, 2019). Read abundances were normalized using the decostand (method = “total”) function of the Vegan package (Okasanen *et al.,* 2019). Beta diversity hypothesis testing was done using Bray-Curtis dissimilarity matrices of the total bacterial community of each sample and subsequently non-metric multidimensional scaling (NMDS). Tests for significance differences in community composition between the substrates, across sampling time point and over the incubation times were performed by analysis of similarity (ANOSIM) and permutation multivariant analysis of variance (PERMANOVA). Additionally, the change in community composition over the course of each incubation was visualized using the average percentage change in abundance of each genus over time (minimum read abundance of 1.0%). This value was calculated by calculating the change in normalised read abundance of each bacterial genus over time compared to the initial community (t0), and highlights only the positive and negative responses of each genus to the substrate additions.

## Acknowledgements

We thank Karl-Peter Rücknagel and Lilly Hufnagel (MPI) for FISH counts of environmental samples during the 2016 Helgoland spring bloom. This study was supported by the Max Planck Society, with additional support by the U.S. National Science Foundation (OCE-1332881 and OCE-1736772 to C.A.) C.A. was also supported by a fellowship from the Hanse Institute for Advanced Study (Delmenhorst, Germany). G.R was also supported by funding from the European Union’s Horizon 2020 research and innovation program under the Marie Sklodowska-Curie grant agreement No. 840804.

## Supplemental Figure Legends

Fig. S1 Diatom abundance and nutrient (silicate, nitrogen, phosphate) concentrations from March 1^st^ to May 30^th^ 2016 at the Helgoland sampling site.

Fig. S2: Bubble plots showing normalized read abundance of members of the *Bacteroidetes* (a), *Gammaproteobacteria* (b), and *Rhodobacteriaceae* (minimum read abundance 0.005%) for each substrate incubation and the unamended incubation at t0, 3d, and 6d for Hel_1 to Hel_4. NMDS plots (right) based on Bray-Curtis dissimilarity reveal separation in *Bacteroidetes*, *Gammaproteobacteria*, and *Rhodobacteriaceae* composition across bloom phases with time. ANOSIM (R:Vegan) indicates significant difference between dissimilarity across bloom phases.

Fig. S3: Shifts in microbial communities (1% minimum abundance) during incubation with FLA-PS. Percentage change in relative read abundance of bacterial genera (and phyla) are shown; the bar chart compares t0 abundances of each genus to 3d and 6d abundances and shows increase or decrease in each genus. ARA = arabinogalactan, CA = carrageenan, CH = chondroitin sulfate, LA = laminarin, XY = xylan, NA = no addition control. (a) Hel_1; note there are no data for CA at 6d (b) Hel_2; Note there are no data for xylan because there are no data from the t0 sample; (c) Hel_3; there are no data for the NA at 6d. Note also that there are no chondroitin data because this substrate was not used at the Hel_3 sample point. (d) Hel_4.

Fig. S4: Relative abundance (% of DAPI stained cells) of *Alteromonas* (black; ALT1413) and *Bacteroidetes* (gray; CF319a) during incubations Hel_1 to Hel_4 enumerated by FISH.

Fig. S5: Percentage of total DAPI-stainable cells at t0 timepoint (∼ 15 minutes after substrate addition) showing staining by laminarin, xylan, chondroitin sulfate, and arabinogalactan for Hel_1 to Hel_4. Note that no chondroitin incubation was carried out for Hel_3. These data are the same as the t0 timepoints in Fig. 6a, replotted here for clarity.

## References

1. Alderkamp, A.-C., van Rijssel, M., and Bolhuis, H. (2007) Characterization of marine bacteria and the activity of their enzyme systems involved in degradation of the algal storage glucan laminarin. FEMS Microbiol Ecol 59: 108–117.

2. Arnosti, C. (2003) Fluorescent derivatization of polysaccharides and carbohydrate-containing biopolymers for measurement of enzyme activities in complex media. J Chromatog B 703: 181–191.

3. Arnosti, C., Reintjes, G., and Amann, R. (2018) A mechanistic microbial underpinning for the size-reactivity continuum of DOC degradation. Mar Chem 206: 93–99.

4. Arnosti, C., Steen, A.D., Ziervogel, K., Ghobrial, S., and Jeffrey, W.H. (2011) Latitudinal gradients in degradation of marine dissolved organic carbon. PLoS ONE 6: e28900.

5. Badur, A.H., Plutz, M.J., Yalamanchili, G., Jagtap, S.S., Schweder, T., Unfried, F. et al. (2017) Exploiting fine-scale genetic and physiological variation of closely related microbes to reveal unknown enzymatic function. J Biol Chem. 292: 13056–13067.

6. Balmonte, J.P., Buckley, A., Hoarfrost, A., Ghobrial, S., Ziervogel, K., Teske, A., and Arnosti, C. (2019) Community structural differences shape microbial responses to high molecular weight organic matter. Environ Microb 21: 557–571.

7. Bauer, M., Kube, M., Telling, H., Richter, M., Lombardot, T., Allers, E. et al. (2006) Whole genome analysis of the marine Bacteroidetes ‘Gramella forsetii’ reveals adaptations to degradation of polymeric organic matter. Environ Microbiol 8: 2201–2213.

8. Behrenfeld, M., Doney, S.C., Lima, I., Boss, E.S., and Siegel, D.A. (2013) Annual cycles of ecological disturbance and recovery underlying the subarctic Atlantic spring plankton bloom. Global Biogeochem Cycles 27: 526–540.

9. Bennke, C.M., Reintjes, G., Schattenhofer, M., Ellrott, A., Wulf, J., Zeder, M., Fuchs, B.M. (2016)Modification of a high-throughput automatic microbial cell enumerations system for shipboard analyses. Appl Environ Microbiol 82: 3289–3296.

10. Boss, E., and Behrenfeld, M. (2010) In situ evaluation of the initiation of the North Atlantic phytoplankton bloom. Geophys Res Lett 37: L18603, doi:10.1029/2010GL044174.

11. Buchan, A., LeCleir, G.R., Gulvik, C.A., and Gonzalez, J.M. (2014) Master recyclers: features and functions of bacteria associated with phytoplankton blooms. Nature Rev Microbio 12: 686–698.

12. Chaplin, M.F., and Kennedy, J.F. (1986) Carbohydrate analysis: A practical approach. IRL Press, Oxford. 228 pp.

13. Cho, K.H., and Salyers, A.A. (2001) Biochemical analysis of interactions between outer membrane proteins that contribute to starch utilization by Bacteroides thetaiotaomicron. J Bact 183: 7224–7230.

14. Cuskin, F., Lowe, E.C., Tample, M.J., Zhu, Y., Cameron, E.A., Pudlo, N.A. et al. (2015) Human gut Bacteroidetes can utilize yeast mannan through a selfish mechanism. Nature 517: 165–173.

15. Daniels, C.J., Poulton, A.J., Esposito, M., Paulsen, M.L., Bellerby, R., St John, M., and Martin, A.P. (2015) Phytoplankton dynamics in contrasting early stage North Atlantic spring blooms: composition, succession, and potential drivers. Biogeosciences 12: 2395–2409.

16. Diepenbroek, M. Glöckner F., Grobe P., Güntsch A., Huber R., König-Ries B., Kostadinov I., Nieschulze J., Seeger B., Tolksdorf R. and Triebel, D. Towards an integrated biodiversity and ecological research data management and archiving platform: The German Federation for the Curation of Biological Data (GFBio) In: Plödereder E, Grunske L, Schneider E, Ull D, editors. Informatik 2014 – Big Data Komplexität meistern. GI-Edition: Lecture Notes in Informatics (LNI) – Proceedings. GI edn. Vol. 232. Bonn: Köllen Verlag; 2014. pp. 1711–1724.

17. Herve, M. (2019) RVAideMemoire: testing and plotting procedures for biostatistics. CRAN https://CRAN.R-project.org/package=RVAideMemoire

18. Hoarfrost, A., Balmonte, J.P., Ghobrial, S., Ziervogel, K., Bane, J., Gawarkiewicz, G., and Arnosti, C. (2019) Gulf Stream ring intrusion on the Mid-Atlantic Bight shelf affects microbially-driven carbon cycling. Frontiers Marine Sci. 6: 394. Doi: 10.3389/fmars.2019.00394

19. Juggins, S. (2016) Rioja: Analysis of quaternary science data. CRAN edition 0.9-21. http://www.staff.ncl.ac.uk/stephen.juggins/

20. Kappelmann, L., Kruger, K., Hehemann, J.-H., Harder, J., Markert, S., Unfried, F. et al. (2018) Polysaccharide utilization loci of North Sea Flavobacteriia as basis for using SusC/D-protein expression for predicting major phytoplankton glycans. The ISME J. 13: 76–91.

21. Kembel, S.W., Ackerly, D.D., Blomberg, S.P., Cornwell, W.K., Cowan, P.D. Helmus, M.R. (2019) ‘picante’ Integrating phylogenies and ecology. CRAN 1.8 https://picante.r-forge.r-project.org

22. Klindworth, A. Pruesse, E. Schweer, T., Peplies J., Quast, C. Horn, M. et al. (2013) Evaluation of general 16S ribosomal RNA gene PCR primers for classical and next-generation sequencing-based diversity studies. Nucleic Acids Res. 41: e1.

23. Krüger, K., Chafee, M., Francis, T.B., delRio, T.G., Becher, D., Schweder, T., et al. (2019) In marine Bacteroidetes the bulk of glycan degradation during algae blooms is mediated by few clades using a restricted set of genes. The ISME J. 13: 2800–2816. doi.org/10.1038/s41396-019-0476-y

24. Landa, M., Blain, S., Harmand, J., Monchy, S., Rapaport, A., and Obernosterer, I. (2018) Major changes in the composition of a Southern Ocean bacterial community in response to diatom-derived dissolved organic matter. FEMS Microb Ecol 94: doi: 10.1093/femsec/fiy034.

25. Landa, M., Cottrell, M.T., Kirchman, D.L., Blain, S., and Obernosterer, I. (2013) Changes in bacterial diversity in response to dissolved organic matter supply in a continuous culture experiment. Aq Microb Ecol 69: 157–168.

26. Li, W., and Godzik, A. (2006) Cd-hit: a fast program for clustering and comparing large sets of protein or nucleotide sequences. Bioinformatics 22: 1658–1659.

27. Mahadevan, A., D’Asaro, E., Lee, C., and Perry, M.J. (2012) Eddy-driven stratification initiates North Atlantic spring phytoplankton blooms. Science 337: 54–58.

28. Manz, W., Amann, R., Ludwig. W., Vancanneyt, M., Schleifer, K.-H.(1996) Application of a suite of 16S rRNA-specific oligonucleotide probes designed to investigate bacteria of the phylum cytophaga-flavobacter-bacteroides in the natural environment. Microbiol 142: 1097–1106.

29. Martens, E.C., Koropatkin, N.M., Smith, T.J., and Gordon, J.I. (2009) Complex glycan catabolism by the human gut microbiota: the Bacteriodetes SUS-like paradigm. J Biol Chem 284: 24673–24677.

30. Martin, A. (2012) The seasonal smorgasbord of the seas. Science 337: 46–47.

31. Martinez, E., Antoine, D., D’Ortenzio, F., and de Boyer Montegut, C. (2011) Phytoplankton spring and fall blooms in the North Atlantic in the 1980s and 2000s. J Geophysical Res 116: C11029, doi:10.1029/2010JC006836.

32. Martinez-Garcia, M., Brazel, D.M., Swan, B.K., Arnosti, C., Chain, P.S.G., Reitenga, K.G. et al. (2012) Capturing single cell genomes of active polysaccharide degraders: an unexpected contribution of Verrucomicrobia. PLoS ONE 7: e35314.

33. Oksanen, J., Blanchet, F.G., Friendly, M., Kindt, R. Legendre, P., McGlinn, D., et al. (2019) vegan: Community Ecology Package. R package. Version 2.5–4. http://CRAN.R-project.org/package=vegan

34. Painter, T.J. (1983) Algal Polysaccharides. In The Polysaccharides. Aspinall, G.O. (ed). New York: Academic Press, pp. 195–285.

35. Pernthaler, A., Pernthaler, J., Amann, R. (2002) Fluorescence in situ hybridization and catalyzed reporter deposition for the identification of marine bacteria. Appl Environ Microbiol 68: 3094–3101.

36. Quast, C., Pruesse, E., Yilmaz, P., Gerken, J., Schweer, T., Yarza, P. et al. (2013) The Silva ribosomal RNA gene database project: Improved data processing and web-based tools. Nucleic Acids Res 41: D590–596.

37. Rakoff-Nahoum, S., Foster, K.R., and Comstock, L.E. (2016) The evolution of cooperation within the gut microbiota. Nature 533: 255–259.

38. Reintjes, G., Arnosti, C., Fuchs, B.M., and Amann, R. (2017) An alternative polysaccharide uptake mechanism of marine bacteria. The ISME J 11: 1640–1650

39. Reintjes, G., Arnosti, C., Fuchs, B.M., and Amann, R. (2019) Selfish, sharing, and scavenging bacteria in the Atlantic Ocean: a biogeographic study of microbial substrate utilisation. The ISME J. 13: 1119–1132. doi.org/10.1038/s41396-018-0326-3

40. Riemann, L., Steward, G.F., and Azam, F. (2000) Dynamics of bacterial community composition and activity during a mesocosm diatom bloom. Appl Environ Microbiol 66: 578–587.

41. RStudio Team (2018) RStudio: integrated development for R. RStudio, Inc. Boston, MA http://www.rstudio.com/.

42. Sarker, S., Feudel, U. Meunier, C., Lemke, P., Dutta, P., and Wiltshire, K.H. (2018) To share or not to share? Phytoplankton species coexistence puzzle in a competition model incorporating multiple resource-limitation and synthesizing unit concepts. Ecol Modelling 383: 150–159.

43. Sarmento, H., and Gasol, J.M. (2012) Use of phytoplankton-derived dissolved organic carbon by different types of bacterioplankton. Environ Microb 14: 2348–2360.

44. Sarmento, H., Morana, C., and Gasol, J.M. (2016) Bacterioplankton niche partitioning in the use of phytoplankton-derived dissolved organic carbon: quantity is more important than quality. The ISME J 10: 2582–2592.

45. Scharfe, M. and Wiltshire, K.H. (2019) Modeling of intra-annual abundance distributions: Constancy and variation in the phenology of marine phytoplankton species over five decades at Helgoland Roads (North Sea). Ecol Modeling 404C: 46–60.

46. Teeling, H., Fuchs, B.M., Becher, D., Klockow, C., Gardebrecht, A., Bennke, C.M. et al. (2012) Substrate-controlled succession of marine bacterioplankton populations induced by a phytoplankton bloom. Science 336: 608–611.

47. Teeling, H., Fuchs, B.M., Bennke, C.M., Kruger, K., Chafee, M., Kappelmann, L. et al. (2016) Recurring patterns in bacterioplankton dynamics during coastal spring algae blooms. eLife 5: e11888. DOI: 10.7554/eLife.11888

48. Teira, E., Gasol, J.M., Aranguren-Gassis, M., Fernandez, A., Gonzalez, J.M., Lekunberri, I., and Alvarez-Salgado, X.A. (2008) Linkages between bacterioplankton community composition, heterotrophic carbon cycling and environmental conditions in a highly dynamic coastal ecosystem. Environ Microb 10: 906–917.

49. Toribio, A., Alako, B., Amid, C., Cerdeno-Tarraga, A., Clarke, L., Cleland, I., et al., (2017) The European nucleotide archive in 2016. Nucleic Acids Res. 45: D36–D40.

50. Unfried, F., Becker, S., Robb, C.S., Hehemann, J.-H., Markert, S., Heiden, S.E. et al. (2018) Adaptive mechanisms that provide competitive advantages to marine bacteroidetes during microalgal blooms. The ISME J. 12: 2894–2906. doi.org/10.1038/s41396-018-0243-5

51. Usov, A.I. (2011) Polysaccharides of the red algae. Adv Carbohydrate Chem Biochem 65: 115–217.

52. Wiltshire, K.H., Boersma, M., Carstens, K., Peters, S., and Scharfe, M. (2015) Control of phytoplankton in a shelf sea: Determination of the main drivers based on the Helgoland Road time series. J Sea Res. 105: 42–52.

53. Xing, P., Hahnke, R.L., Unfried, F., Markert, S., Hugang, S., Barbeyron, T. et al. (2014) Niches of two polysaccharide-degrading Polaribacter isolates from the North Sea during a spring diatom bloom. The ISME J. 9: 1410–1422.

54. Yilmaz, P. Kottmann, R. Field, D., Knight, R., Cole, J.R., Amaral-Zettler, L., et al. (2011) Minimum information about a marker gene sequence (MIMARKS) and minimum information about any (x) sequence (MIxS) specifications. Nat Biotech 29: 415–420.

55. Zimmerman, A.E., Martiny, A.C., and Allison, S.D. (2013) Microdiversity of extracellular enzyme genes among sequenced prokaryotic genomes. The ISME J. 7: 1187–1199.

