## Supplemental Information for "Short-term changes in polysaccharide utilization mechanisms of marine bacterioplankton during a spring phytoplankton bloom"

Supplemental Figure Legends:

Fig. S1 Diatom abundance and nutrient (silicate, nitrogen, phosphate) concentrations from March 1$^{st}$ to May 30$^{th}$ 2016 at the Helgoland sampling site.

Fig. S2: Bubble plots showing normalized read abundance of members of the *Bacteroidetes* (a), *Gammaproteobacteria* (b), and *Rhodobacteriaceae* (minimum read abundance 0.005%) for each substrate incubation and the unamended incubation at t0, 3d, and 6d for Hel_1 to Hel_4. NMDS plots (right) based on Bray-Curtis dissimilarity reveal separation in *Bacteroidetes*, *Gammaproteobacteria*, and *Rhodobacteriaceae* composition across bloom phases with time. ANOSIM (R:Vegan) indicates significant difference between dissimilarity across bloom phases.

Fig. S3: Shifts in microbial communities (1% minimum abundance) during incubation with FLA-PS. Percentage change in relative read abundance of bacterial genera (and phyla) are shown; the bar chart compares t0 abundances of each genus to 3d and 6d abundances and shows increase or decrease in each genus. ARA = arabinogalactan, CA = carrageenan, CH = chondroitin sulfate, LA = laminarin, XY = xylan, NA = no addition control. (a) Hel_1; note there are no data for CA at 6d (b) Hel_2; Note there are no data for xylan because there are no data from the t0 sample; (c) Hel_3; there are no data for the NA at 6d. Note also that there are no chondroitin data because this substrate was not used at the Hel_3 sample point. (d) Hel_4.

Fig. S4: Relative abundance (% of DAPI stained cells) of *Alteromonas* (black; ALT1413) and *Bacteroidetes* (gray; CF319a) during incubations Hel_1 to Hel_4 enumerated by FISH.

Fig. S5: Percentage of total DAPI-stainable cells at t0 timepoint (~ 15 minutes after substrate addition) showing staining by laminarin, xylan, chondroitin sulfate, and arabinogalactan for Hel_1 to Hel_4. Note that no chondroitin incubation was carried out for Hel_3. These data are the same as the t0 timepoints in Fig. 6a, replotted here for clarity.

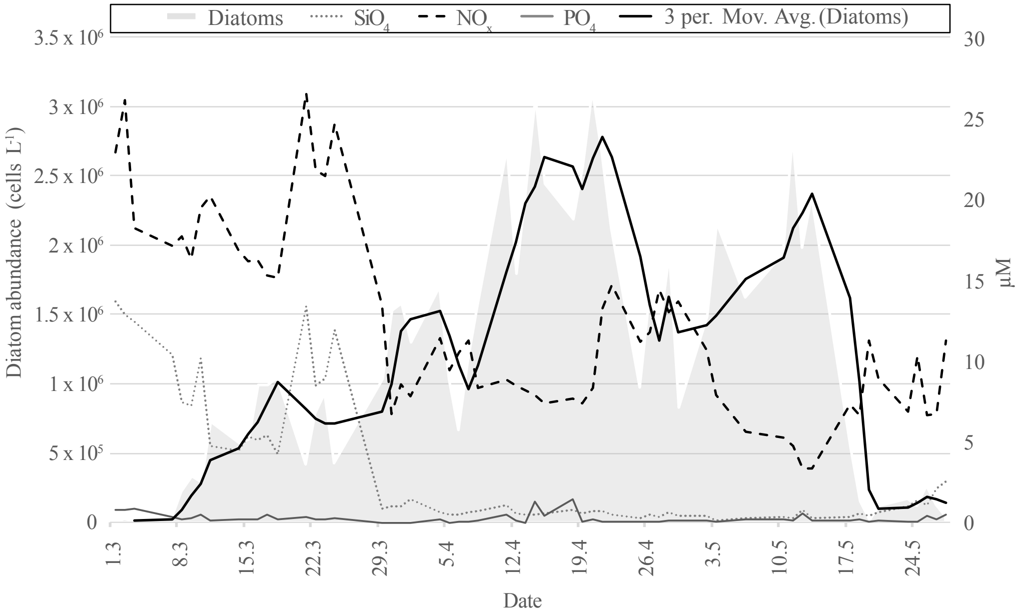

Diatoms
$SiO_4$
$NO_x$
$PO_4$
3 per. Mov. Avg. (Diatoms)
3.5 x $10^6$
3 x $10^6$
2.5 x $10^6$
2 x $10^6$
1.5 x $10^6$
1 x $10^6$
5 x $10^5$
0
Diatom abundance (cells $L^{-1}$)
30
25
20
15
10
5
0
µM
1.3
8.3
15.3
22.3
29.3
5.4
12.4
19.4
26.4
3.5
10.5
17.5
24.5
Date

### a) Bacteroidetes

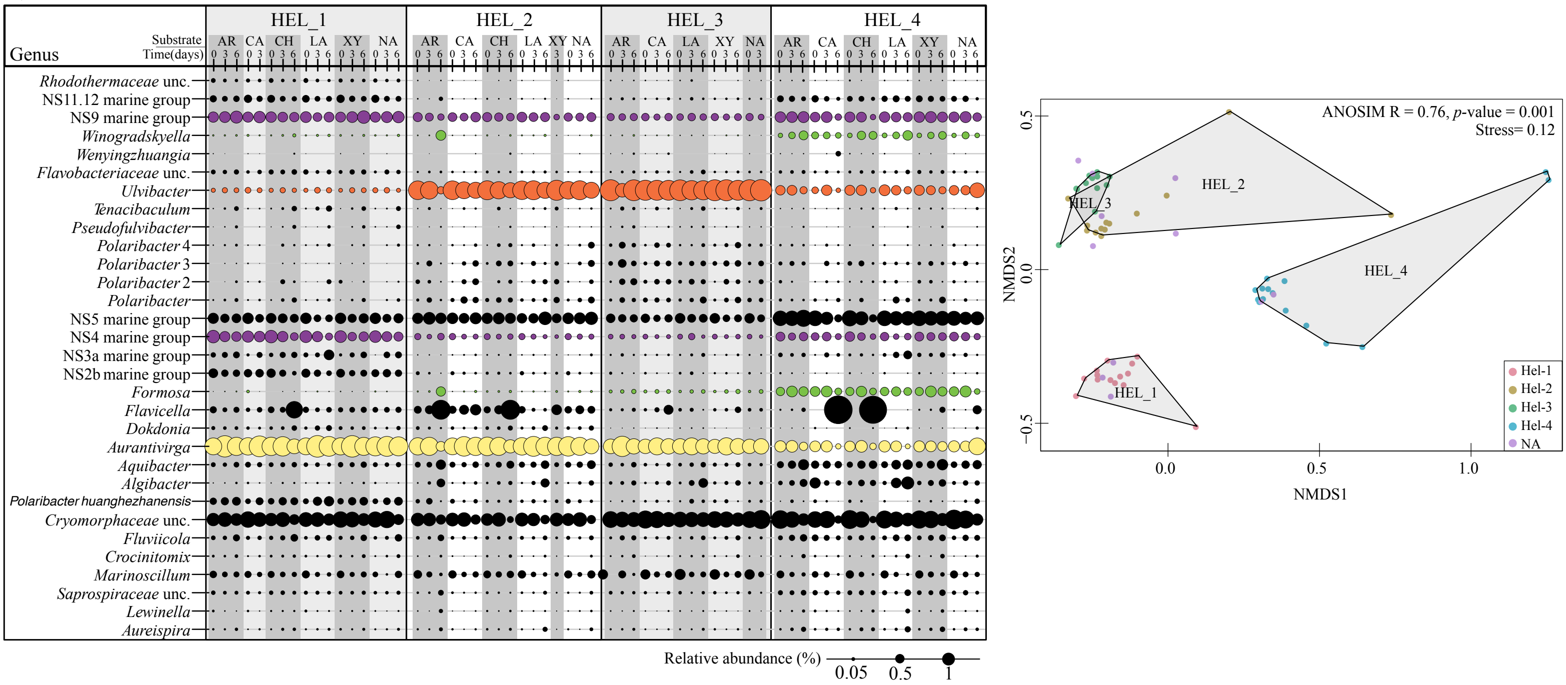


### b) Gammaproteobacteria

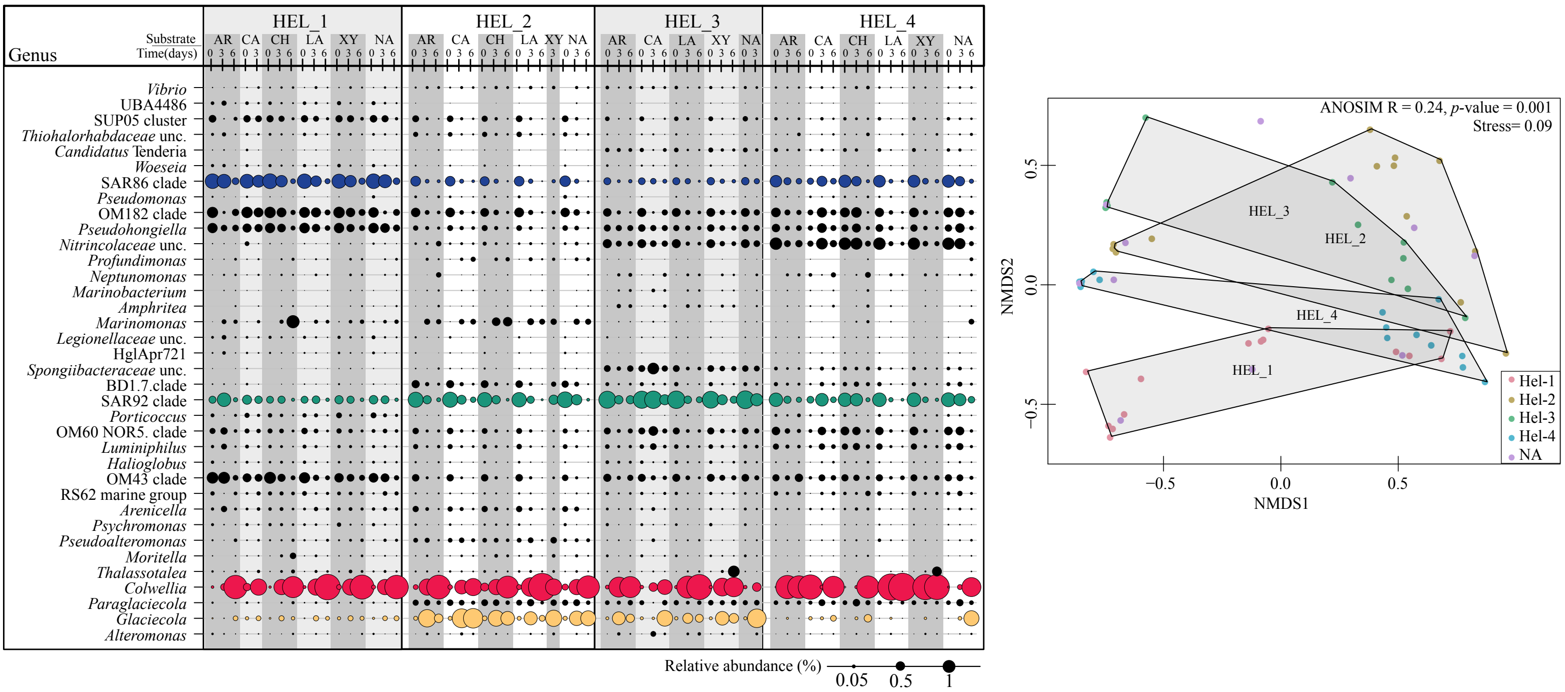


### c) Rhodobacteraceae

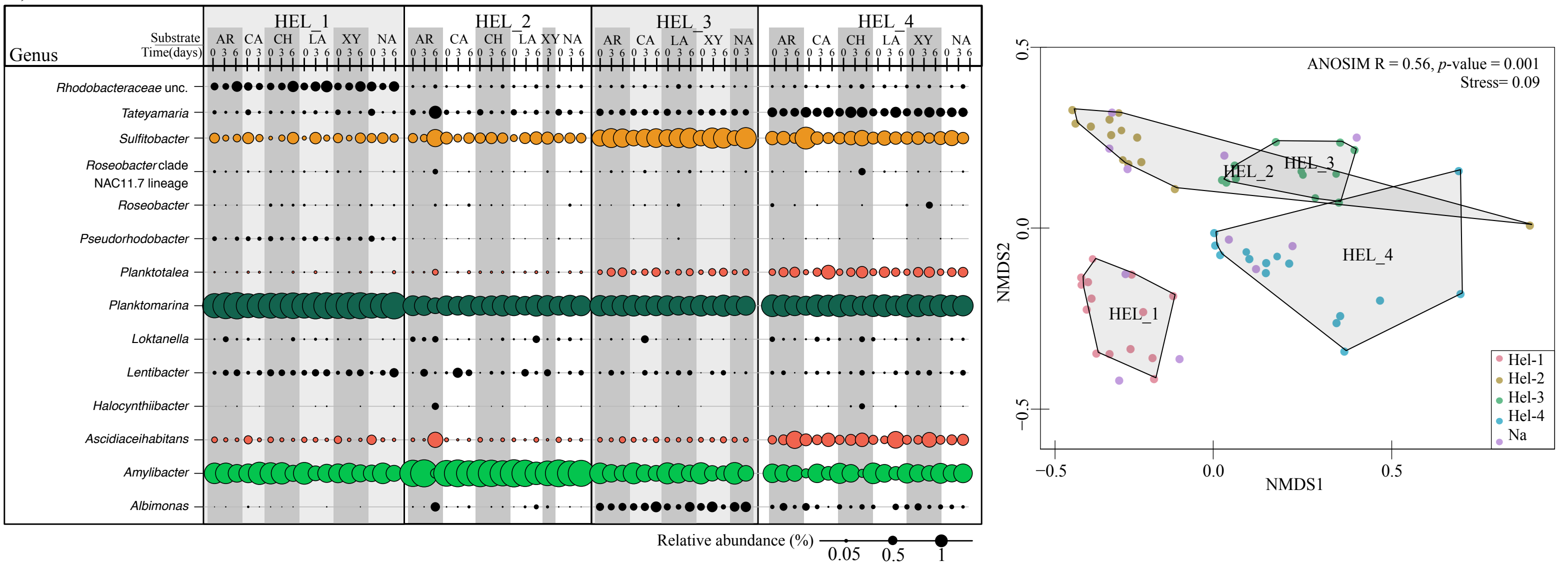


Reintjes et al. Supplementary Figure S2

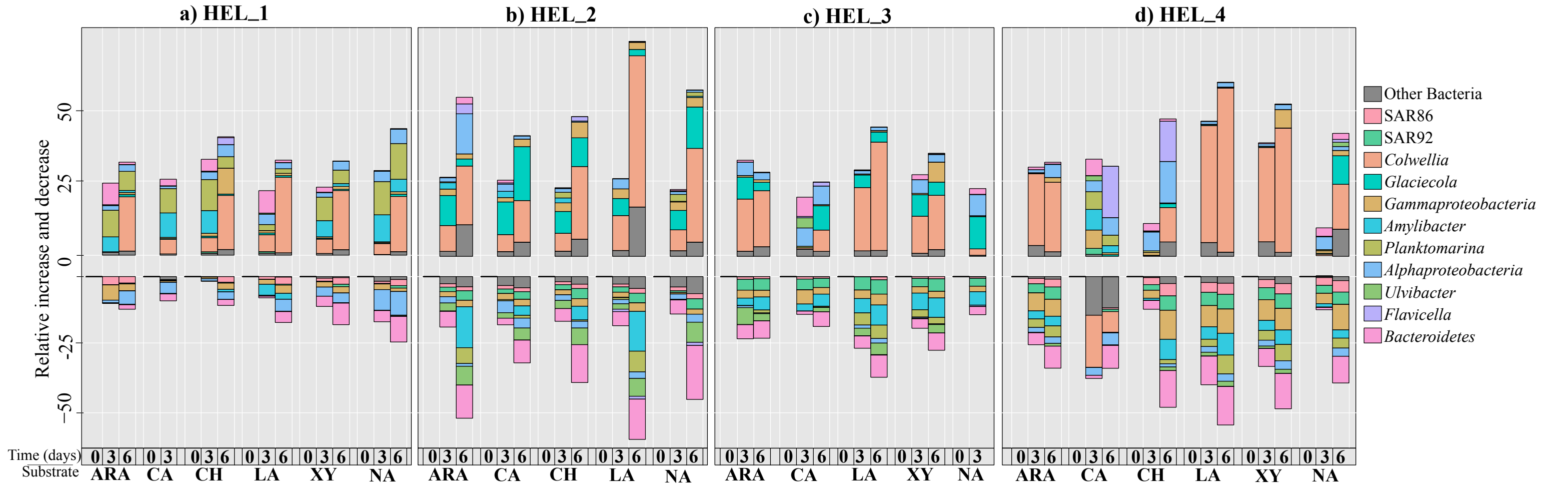


Reintjes et al. Supplementary Figure S3

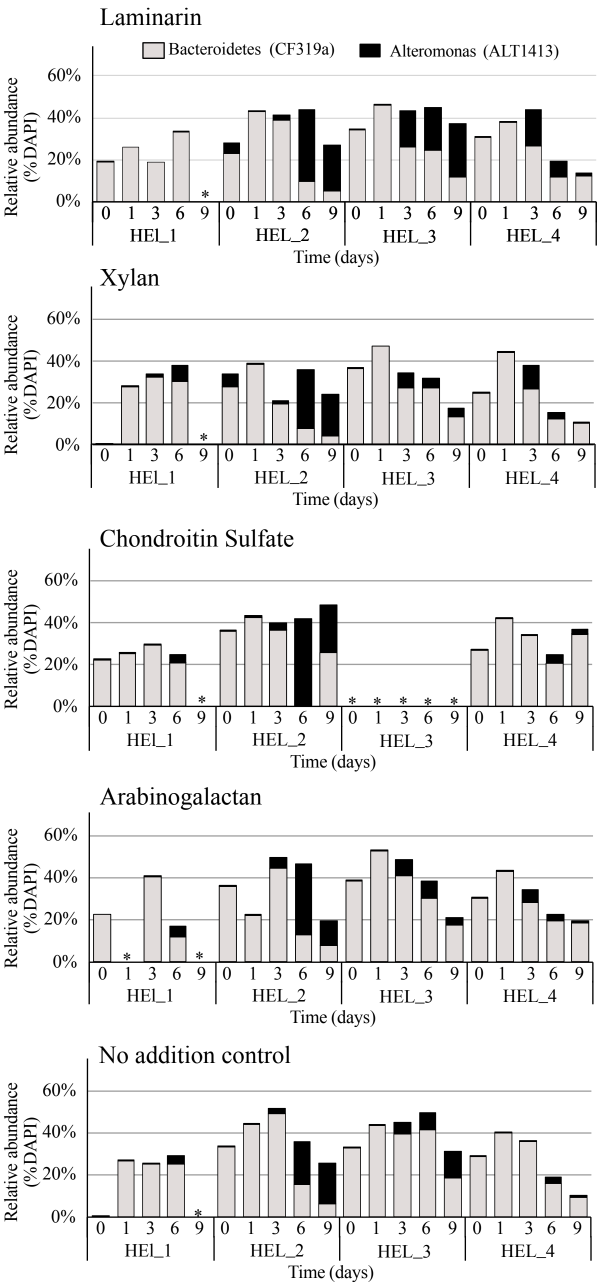


Reintjes et al. Supplementary Figure S4

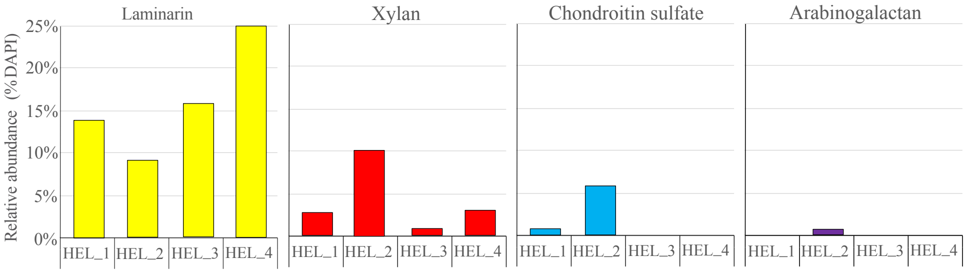


Reintjes et al. Supplementary Figure S5
